# Two-dimensional reward evaluation in mice

**DOI:** 10.1101/2020.04.14.040808

**Authors:** Vladislav Nachev, Marion Rivalan, York Winter

## Abstract

When choosing among multi-attribute options, integrating the full information may be computationally costly and time-consuming. So-called non-compensatory decision rules only rely on partial information, for example when a difference on a single attribute overrides all others. Such rules may be ecologically more advantageous, despite being economically suboptimal. Here we present a study that investigates to what extent animals rely on integrative rules (using the full information) versus non-compensatory rules when choosing where to forage. Groups of mice were trained to obtain water from dispensers varying along two reward dimensions: volume and probability. The mice’s choices over the course of the experiment suggested an initial reliance on integrative rules, later displaced by a sequential rule, in which volume was evaluated before probability. Our results also demonstrate that while the evaluation of probability differences may depend on the reward volumes, the evaluation of volume differences is seemingly unaffected by the reward probabilities.

## Introduction

Animals confronted with options that differ on a single attribute generally make economically rational choices consistent with gain maximization (Monteiro, Vasconcelos, and Kacelnik 2013; Rivalan, Winter, and Nachev 2017). In multi-attribute choice (Pitz and Sachs 1984; Jansen, Duijvenvoorde, and Huizenga 2012; Hunt, Dolan, and Behrens 2014) however, where reward attributes must be weighed against each other (price *vs.* quality, risk *vs.* payoff, etc.), consistent deviations from economical rationality have been described in humans (Tversky and Kahneman 1974; Rieskamp, Busemeyer, and Mellers 2006; Katsikopoulos and Gigerenzer 2008) and non-human animals (Shafir, Waite, and Smith 2002; Bateson, Healy, and Hurly 2003; Schuck-Paim, Pompilio, and Kacelnik 2004; Scarpi 2011; Nachev and Winter 2012; Nachev et al. 2017; Constantinople, Piet, and Brody 2019). Some deviations from gain maximization can be accounted for by considering the ecological circumstances of an animal, which may confer fitness benefits to seemingly irrational choices (Kacelnik 2006; Houston, McNamara, and Steer 2007; Trimmer 2013; McNamara, Trimmer, and Houston 2014).

An animal foraging in its natural environment mostly encounters food items that differ on multiple attributes, but only some of those attributes affect the long-term gains. We refer to those attributes as reward dimensions. In multidimensional choice the decision task is considerably simplified if differences that are (nearly) equal are not evaluated but ignored (Tversky 1969; Pitz and Sachs 1984; Shafir 1994; Shafir and Yehonatan 2014). For example, an animal might only consider the one reward dimension (e.g. prey size) that most strongly affects the long-term gains. Such decision processes in which one reward dimension overrides the others have been described as non-compensatory (Pitz and Sachs 1984; Reid et al. 2015) and can potentially increase speed of decision and decrease computation costs at the expense of accuracy. Attributes can be considered sequentially, for example ranked by salience, until a sufficient difference is detected on one attribute so that a decision can be reached (Brandstätter, Gigerenzer, and Hertwig 2006; Jansen, Duijvenvoorde, and Huizenga 2012). In compensatory decision-making (Pitz and Sachs 1984; Reid et al. 2015) on the other hand, choice is affected by multiple attributes that are integrated into a common decision currency (utility) (Levy and Glimcher 2012). A fully integrative approach that makes use of all the available information (also referred to as absolute reward evaluation Tversky 1969; Shafir 1994; Shafir and Yehonatan 2014) is equivalent to gain maximization. For example, if options differ along the reward dimensions of amount and probability of obtaining this amount, maximizing the gain is ensured by selecting the option with the highest expected value, which is the product of the amount and probability. Even in two-dimensional reward evaluation, a range of strategies are possible, from sequential and other non-compensatory rules, up to full integration.

When studying animal decision-making, preferences are measured over many choices, especially when options differ in reward probability. Although a rational subject should exclusively select the most profitable option, animals can persist in choosing less profitable options even after long training, usually at some low frequency (Kacelnik 1984). The partial preference observed in choice experiments can be explained by profitability matching (Kacelnik 1984), which states that animals proportionally allocate their effort depending on the relative pay-off of the options.

Scalar Utility Theory (SUT: Kacelnik and Brito e Abreu 1998; Marsh and Kacelnik 2002) is a framework that proposes a proximate mechanism that accounts for partial preferences in the context of reward amount and reward variability (Rosenström, Wiesner, and Houston 2016). Based on findings in psychophysics, SUT postulates that cognitive representations of stimuli exhibit a scalar property, i.e. they have error distributions that are normal with a mean equal to the magnitude of the stimulus and a standard deviation that is proportional to the mean. In other words, SUT states that the memory traces of perceived or expected outcomes of choices are subject to Weber’s Law (Akre and Johnsen 2014) and that rewards are evaluated proportionally rather than linearly (Marsh and Kacelnik 2002; Rosenström, Wiesner, and Houston 2016). Therefore, according to SUT choice is modelled by sampling from the internal representations of the choice options and selecting the most favorable sample. This allows for making quantitative predictions about the strength of preferences from the contrasts between options.

In previous experiments we have demonstrated that proportional processing can be used to predict the choice behavior of animals when options vary along a single dimension (Nachev, Stich, and Winter 2013; Rivalan, Winter, and Nachev 2017). In the present study we extend the application of proportional processing and SUT to two-dimensional choice tasks with the aim to test whether (contradictory) information from two reward dimensions generates choices more consistent with integrative or non-compensatory decision rules. We used a combination of behavioral studies of mice and a decision-making model based on SUT.

## Animals, Methods, and Materials

### Animals

The experiments were conducted with three cohorts of C57BL/6NCrl female mice (Charles River, Sulzfeld, Germany, total n = 30). Mice were five weeks old on arrival. The mice from each cohort were housed together, before and during the experiments. They were marked with unique radiofrequency identification tags (RFID: 12 × 2.1 mm, 125 kHz, Sokymat, Rastede, Germany) under the skin in the scruff of the neck and also earmarked at age six weeks. At age seven weeks mice were transferred to the automated group home cage for the main experiment. Pellet chow (V1535, maintenance food, ssniff, Soest, Germany) was always accessible from a trough in the cage lid. Water was available from the operant modules of the automated group cage, depending on individual reward schedules. Light conditions in the experiments were 12:12 LD and climatic conditions were 23 ± 2°C and 50–70% humidity.

#### Ethics statement

The experimental procedures were aimed at maximizing animal welfare. During experiments, mice remained undisturbed in their home cage. Data collection was automated, with animals voluntarily visiting water dispensers to drink. The water intake and health of the mice was monitored daily. Due to the observational nature of the study, animals were free from damage, pain, and suffering. The animals were not sacrificed at the end of the study, which was performed under the supervision and with the approval of the animal welfare officer heading the animal welfare committee at Humboldt University. Experiments followed national regulations in accordance with the European Communities Council Directive 10/63/EU.

### Cage and dispenser system

We used two automated home cages (612 × 435 × 216 mm, P2000, Tecniplast, Buggugiate, Italy) with woodchip bedding (AB 6, AsBe-wood, Gransee, Germany), and enriched with two grey PVC tubes and paper towels as nesting material. The cage was outfitted with four computer-controlled liquid dispensers. The experimental set-up of cage 1 is described in detail in Rivalan, Winter, and Nachev (2017). Briefly, mice were detected at the dispensers via infrared beam-break sensors and RFID-sensors. Water delivery at each dispenser could be controlled, so that it could be restricted or dispensed at different amounts on an individual basis. Mice were therefore rewarded with droplets of water from the dispenser spout that they could remove by licking. We changed cage bedding and weighed all animals on a weekly basis, always during the light phase and at least an hour before the start of the testing session. Data were recorded and stored automatically on a laptop computer using PhenoSoft Control software (PhenoSys, Germany). Time-stamped nose poke events and amounts of water delivered were recorded for each dispenser, with the corresponding mouse identity.

The second automated group cage (cage 2) was made for the purposes of this study and was nearly identical to cage 1. The crucial modification was that the stepping-motor syringe pump was replaced with a model that used disposable plastic 25-mL syringes (cage 2) instead of gas-tight Hamilton glass syringes (Series 1025, cage 1). Thus, the pumping systems in the two cages differed in the smallest reward that could be delivered and in the precision of reward delivery (mean ± SD: 0.33 ± 0.03 *μLstep^−^*^1^ in cage 1 vs. 1.56 ± 0.24 *μLstep^−^*^1^ in cage 2). The precision of each pump was estimated by manually triggering reward visits at different preset pump steps (17 and 42 in cage 1, 3 and 12 in cage 2) and collecting the expelled liquid in a graduated glass pipette placed horizontally next to the cage. Each dispenser was measured by the same trained experimenter at least 20 times for each pump step value.

### Experimental schedule

The general experimental procedure was as described before (Rivalan, Winter, and Nachev 2017). The water dispensers were only active during an 18h-long drinking session each day that began with the onset of the dark phase and ended six hours after the end of the dark phase. The reward properties (volume and probability) were dependent on the experimental condition. Rewards were drawn from fixed pseudo-random repeating sequences. These sequences were: 11101111101101111110 for 80%, 11011101110101101110 for 70%, 10110101101001001010 for 50%, 10010100100001001000 for 30%, and 10001000010001000000 for 20%, where 1 is a rewarded nose poke and 0 is an unrewarded nose poke.

Although individual mice shared the same dispensers inside the same cage, they were not necessarily in the same experimental phase during training or experimental condition in the main experimental phase. The three cohorts (1-3 in chronological order) were tested consecutively, with cohort 2 housed in cage 2 and the other cohorts housed in cage 1. If after any drinking session during any experimental phase a mouse drank less than 1 *mL* of water, we placed two water bottles in the automated cage, gently awakened all mice and allowed them to drink freely until they voluntarily stopped. An overview of the training and experimental phases is given in Figure 1a.

**Figure 1:**
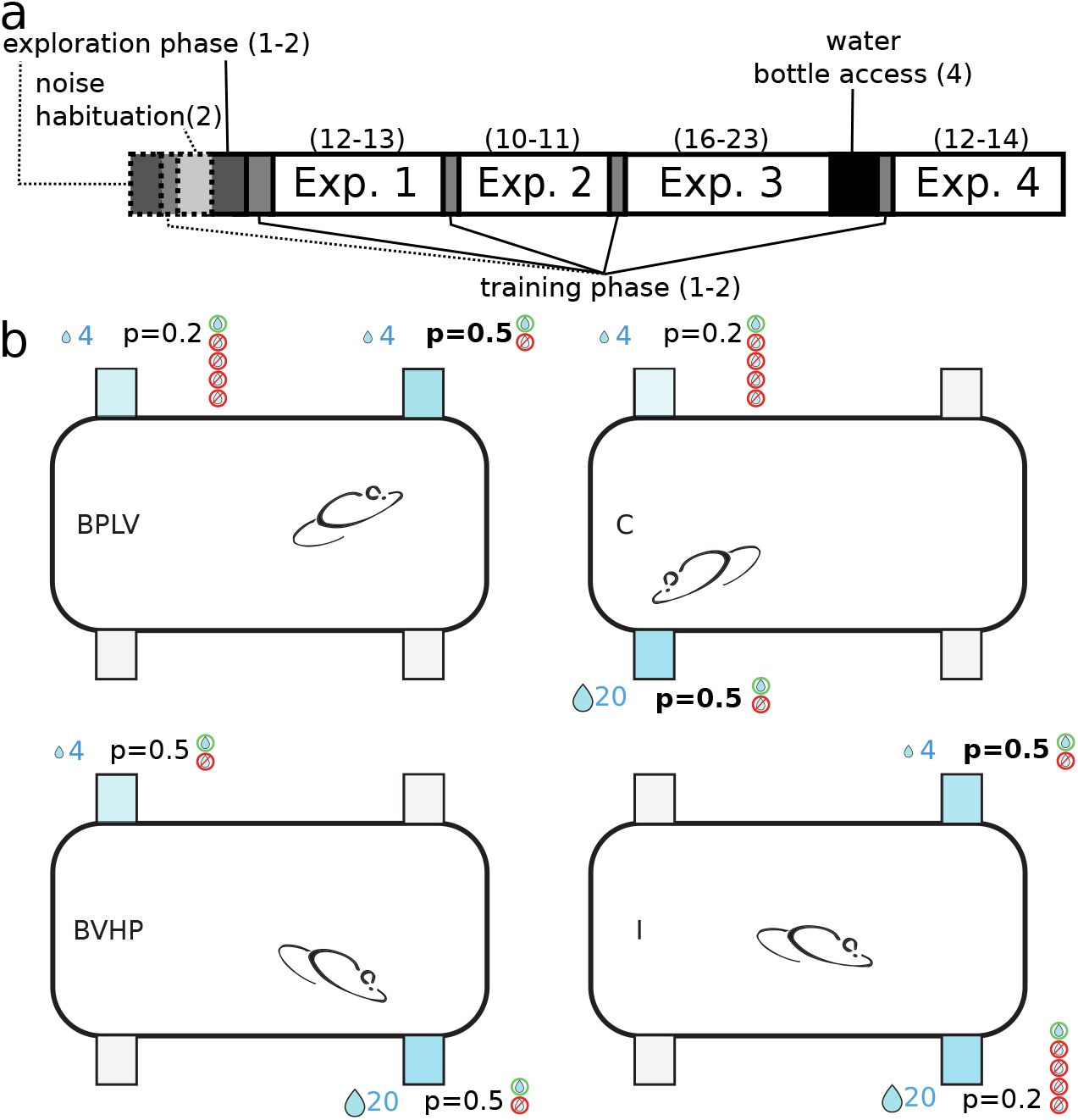
Experimental conditions and schedules. (**a**) Experimental schedule with all phases. The number of days are given in parentheses. Mice began with an exploration phase, followed by a training phase. Before every experiment (Exp. 1-4) there was another training phase for one day. Between experiments 3 and 4 there was a four-day break with water from a bottle. Phases shown with dashed lines were only present in the schedule for cohort 3, because four mice had difficulties advancing beyond the exploration phase. (**b**) Behavioral task in four conditions (BPLV, BVHP, C, and I) of experiment 1. Mice were free to nosepoke in all four corners, two of which (shown in blue) were rewarding for the example mouse, with the reward properties shown above or below the reward corners. For clarity, only one example mouse of the eight mice is shown, with other mice experiencing different conditions at different dispensers. BPLV: baseline for probability at low volume; BVHP: baseline for volume at high probability; C: congruent condition; I: incongruent condition.

#### Exploration phase

At the beginning of this phase there were ten mice in each cohort, except for cohort 2, in which one mouse was excluded due to the loss of the RFID tag after implantation (the mouse was in good health condition). The mice were transferred to the automated cages 1-2 hours before the first drinking session of the exploration phase. The purpose of this phase was to let mice accustom to the cage and learn to use the dispensers to obtain water. Therefore, each nose poke at any dispenser was rewarded with a constant volume of 20 *μL*. The criterion for advancing to the following training phase was consuming more than 1 *mL* in a single drinking session. Mice that did not reach the criterion remained in the exploration phase until they either advanced to the following phase or were excluded from the experiment (*n* = 1 mouse in cohort 2).

#### Training phase

In this phase the reward volume was reduced to 10 *μL* and the reward probability was reduced to 0.3 at all dispensers. These reward values ensured that mice remained motivated to make several hundred visits per drinking session. Associative learning is also enhanced by the unpredictability of the expected reinforcer (Maddux et al. 2007). The training phase was repeated for one to two days until at least eight mice fulfilled the criterion of consuming more than 1 *mL* of water in one drinking session. The purpose of the training phase was to introduce mice to the reward dimensions (volume and probability) that would be used in the following discrimination experiments. In cohorts 1 and 2, mice were excluded from the experiment if they did not reach the criterion in two days, or, alternatively, if more than eight mice had reached the criterion, mice were excluded at random to ensure a balanced number of mice per dispenser. These mice were returned to regular housing.

#### Noise habituation

We introduced a noise habituation phase for the mice in cohort 3, because after two days only six of them had advanced to the training phase (Fig. 1a). The unusually low number of visits made by mice that did not pass the exploration phase suggested that the noise produced by the pumping systems might scare naive, shy mice away from the dispensers. In order to ensure that all mice were successfully trained, we designed the noise habituation so that rewards at all four dispensers were delivered at regular intervals (7 *μL* every minute), regardless of the behavior of the mice. After two days, all mice had made at least 200 nose pokes and the cohort then continued with either exploration or training. Two days later all mice successfully completed the training phase and two mice were randomly selected for removal from the experiment, bringing the number of mice to eight. We therefore updated our training procedure to always begin with noise habituation, followed by the exploration phase and the training phase.

### General procedure in the main experiments

After eight mice had successfully passed the training phase, they proceeded with experiment 1 from the main experiments (1-4). In all of the main experiments mice had a choice between four dispensers, where two were not rewarding and the other two gave rewards with volumes and probabilities that depended on the experimental condition (Figs. 1b, 2). In most conditions one of the rewarding dispensers (high-profitability dispenser) was more profitable than the other (low-profitability dispenser). The sequence of conditions was randomized for each individual, so that any given mouse was usually experiencing a different experimental condition than all other mice. On any given day two of the dispensers were rewarding for four mice and the other two were rewarding for the other four mice. Within each group of four, each pair of mice shared the same high and low-profitability dispensers, which were spatially inverted between pairs of mice. This pairing was done to increase the throughput of the experiments, while controlling for potential social learning effects and distributing mice evenly over the dispensers to minimize crowding effects.

The behavioral measure of interest was the relative visitation rate to the high-profitability dispenser that could develop in one drinking session. Choice behavior in sequential testing with multiple conditions can be influenced by the previous conditions and by side bias. We aimed to mitigate the sequential effects through randomization, and the side bias through spatially reversing the choice options. As a control for positional biases, each condition was followed by a reversal on the next day, so that the high and low-profitability dispensers were spatially inverted for all mice, whereas the two non-rewarding dispensers remained unchanged. Reversal was followed by the next experimental condition, with pseudo-random distribution of the dispensers among the pairs of mice following the constraints described above. The reversal condition is potentially harder to learn and may represent the lower bound of choice performance, but its exclusion from the results did not lead to any qualitative changes. Over the 50 total days in the main experiment (twice the number of conditions shown in Fig. 2, because of reversals, plus experiment 4), each mouse experienced each dispenser as a high-profitability dispenser between 11 and 14 times. In the event of an electrical or mechanical malfunction, data from the failed condition and its reversal were discarded and the failed condition was repeated at the end of the experiment, lengthening slightly the duration of the experiment. Such a failure occurred once in cohort 1, four times in cohort 2 and did not occur in cohort 3. After experiments 1 and 2, mice were given another training phase (rewards with 10 *μL* and 0.3 probability) for a single day, before they proceeded with the next experiment. After experiment 3 mice were given water from a standard water bottle for four days (with water dispensers inactive), followed by one day in the training phase, before proceeding with experiment 4. At the end of experiment 4 mice were returned to the animal facility.

**Figure 2:**
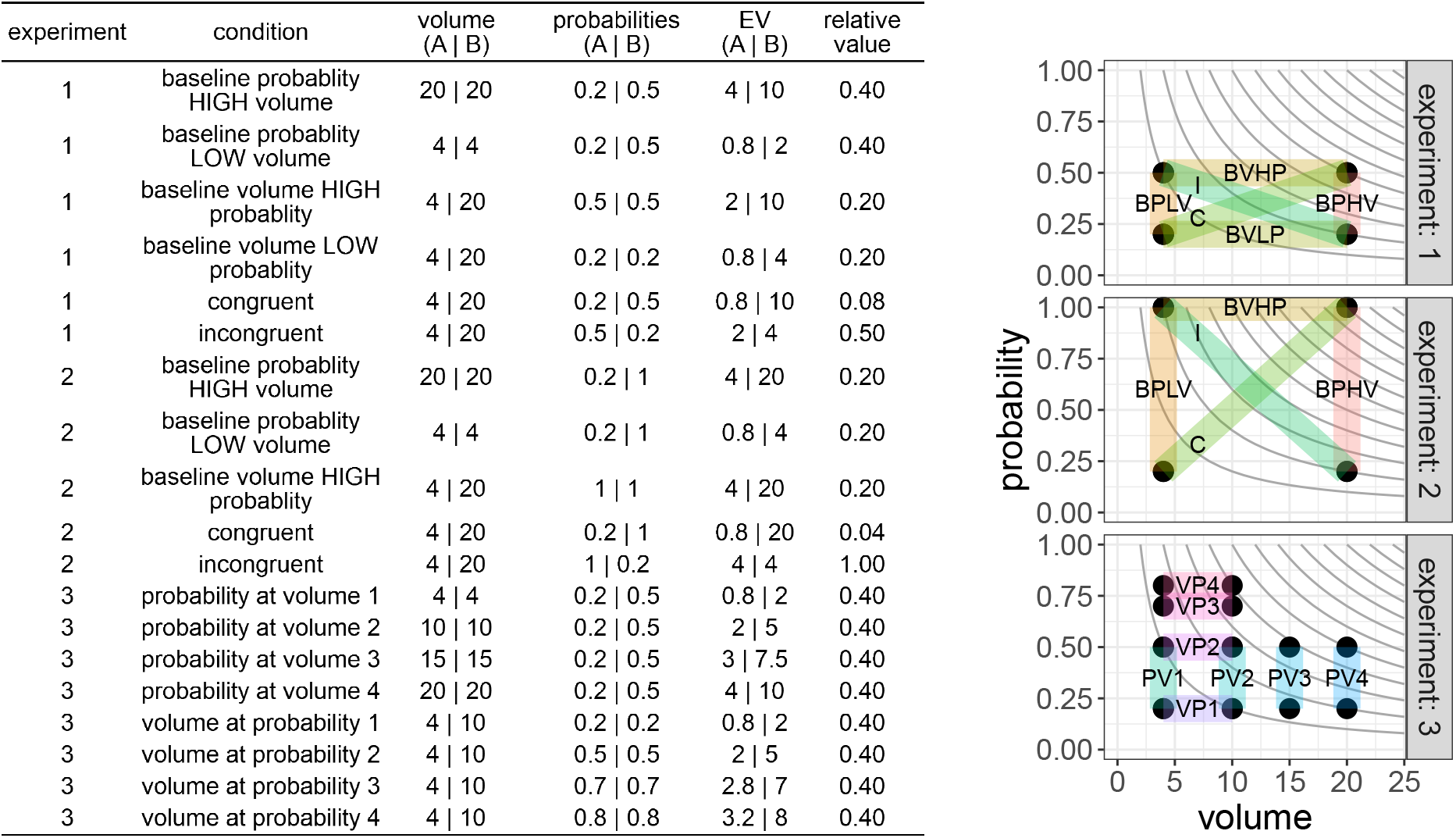
Overview of the experimental conditions in all four experiments. Options (A and B) differed on one or both reward dimensions (reward volume and probability), resulting in different expected values (EV). The black dots give the volume and probability for each option. The transparent segments connect the two options available in each condition. Gray curves give points of equal expected value (EV = volume × probability). The relative value is *EV_A_/EV_B_*. The conditions in experiment 4 were identical to those in experiment 1. The baseline for volume at low probability condition (BVLP) in experiment 1 was not repeated in experiment 2, but instead the results from experiment 1 were reused in further analyses. Condition sequences were randomized for each mouse. Volumes shown (in *μL*) are for cohorts 1 and 3. In cohort 2 the volumes were 4.7 instead of 4, 9.4 instead of 10, 14.0 instead of 15, and 20.3 instead of 20 *μL*.

#### Experiment 1

In the baseline conditions rewards only differed on one reward dimension (the relevant dimension), but not on the other dimension (the background dimension). For example, in the baseline for probability at low volume (BPLV) condition, both options had the same volume of 4 *μ*L, but one option had a probability of 0.2 and the other, a probability of 0.5 (Figs. 1b, 2). In the baseline for volume conditions (Fig. 1b) both rewarding options had the same probabilities (either 0.2, baseline for volume at low probability, BVLP; or 0.5, baseline for volume at high probability, BVHP), but one had a volume of 4 *μ*L, and the other had a volume of 20 *μ*L. Based on previous experiments (Rivalan, Winter, and Nachev 2017), we expected a baseline difference between 4 *μ*L and 20 *μ*L volumes to result in a similar discrimination performance (relative preference for the superior option) compared to a baseline difference between probabilities 0.2 and 0.5. In the congruent (C) condition one option was superior to the other on both dimensions (Fig. 1b). Finally, in the incongruent (I) condition each of the options was superior to the other on one of the reward dimensions, so that the option that had the higher volume had the lower probability and vice versa (Fig. 1b). The main goals of this experiment were to (1) test whether the baseline performance when only one dimension was relevant was a good predictor for the discrimination performance in the congruent and incongruent conditions when both dimensions were relevant and (2) whether the trade-off between dimensions affected preference in the incongruent condition. Since the differences on both dimensions were chosen to be of comparable salience (Rivalan, Winter, and Nachev 2017), we expected the mean discrimination performance in the incongruent condition to be at chance level (0.5), despite the difference in expected value (Fig. 2).

#### Experiment 2

In previous experiments (Rivalan, Winter, and Nachev 2017) we had shown that the relative stimulus intensity (*i*), i.e. the absolute difference between two options divided by their mean (difference/mean ratio), was a good predictor of discrimination performance for both volume and probability differences. Another finding from these experiments was that, at least initially, mice responded less strongly to differences in volume than to differences in probability, despite equivalence in expected values (Rivalan, Winter, and Nachev 2017). We aimed to correct for this effect in experiment 1 by selecting options with a higher relative intensity for volume (4 *μ*L vs. 20 *μ*L, *i* = 1.33) than for probability (0.2 vs. 0.5, *i* = 0.857). In experiment 2 we wanted to test whether mice would exhibit a decreased sensitivity for volume when both reward dimensions had the same relative intensity (*i* = 1.33). Thus, for the conditions in experiment 2 we simply replaced the 0.5 probability from the conditions in experiment 1 with a probability of 1 (Fig. 2). We did not repeat the BVLP condition, in which both probabilities were set at 0.2. With the two choice options having the same expected values, we hypothesized that the discrimination performance in the incongruent condition would be at chance level if both dimensions were equally weighed and equally perceived. On the other hand, if mice were less sensitive for volume than for probability differences as in our previous experiments, then the discrimination performance in the incongruent condition should be skewed towards probability (< 0.5).

#### Experiment 3

In the previous experiments we used two different baseline conditions for each dimension (BPLV, BPHV, BVLP, and BVHP, Fig. 2), in order to exhaust all combinations of reward stimuli and balance the experimental design. However, we also wanted to test whether the level of the background dimension despite being the same across choice options nevertheless affected the discrimination performance on the relevant dimension. If mice use a non-compensatory decision rule, we can predict that regardless of the level of the background dimension, the discrimination performance on the relevant dimension should remain constant. Alternatively, with absolute reward evaluation the subjective difference between the options is said to decrease as the background (irrelevant) dimension increases and therefore the discrimination performance is also expected to decrease (Shafir and Yehonatan 2014). This prediction is derived from the concave shape of the utility function, which is generally assumed to increase at a decreasing rate with the increase in any reward dimension (Kahneman and Tversky 1979; Kenrick et al. 2009; but see also Kacelnik and Brito e Abreu 1998). The same prediction can be made if we assume that motivation decreases with satiety, i.e. the strength of preference decreases under rich environmental conditions (Schuck-Paim, Pompilio, and Kacelnik 2004), for example at high reward volume or probability. In order to test whether the two reward dimensions (volume and probability) interact with each other even when one of them is irrelevant (as background dimension that is the same across choice options), we performed experiment 3.

The conditions in experiment 3 were chosen to be similar to the baseline conditions in the previous experiments, by having one background and one relevant dimension (Fig. 2). The relevant dimension always differed between the two options. For the probability dimension, we selected the same values of 0.2 and 0.5 (*i* = 0.86), as in the previous experiments. For the volume dimension we selected the values of 4 *μ*L and 10 *μ*L (4.8 *μ*L and 9.6 *μ*L in cohort 2, Fig. 2), because these values have the same relative intensity as the two probabilities. Furthermore, the combination of a higher volume with a probability of 0.8 was expected to result in an insufficient number of visits for analysis. Cohort 2 had different reward volumes due to differences in the pumping process between the two cages used (Cage and dispenser system), which also resulted in a lower relative intensity for volume (*i* = 0.67 instead of 0.86; we will return to this point in the discussion). There were four different levels for each background dimension (volume and probability, Fig. 2). Each mouse had its own pseudo-random sequence of the eight possible conditions.

#### Experiment 4

For laboratory mice that have unrestricted access to a water bottle, the volume of a water reward is not usually a stimulus that predicts reward profitability. In previous experiments (Rivalan, Winter, and Nachev 2017), mice had shown an improved discrimination performance for volume over time. This suggests that with experience mice become more attuned to the relevant reward dimension. In order to test whether the discrimination performance for one or both dimensions improved over time, we performed experiment 4, which had the same conditions as experiment 1 (Fig. 2), but with a new pseudo-random order. The same mice participated in all experiments (1 to 4), with about seven weeks between experiment 1 and experiment 4.

### Data analysis

Data analysis and simulations were done using R (Team 2020). All data and code are available in the Zenodo repository: https://doi.org/10.5281/zenodo.4223729.

On average (mean ± SD), mice made 477 ± 163 nose pokes per drinking session (Fig. S1), with a mean proportion of 0.79 ± 0.1 nose pokes at the rewarding dispensers. In order to analyze choices only after mice had some experience with each option (Rivalan, Winter, and Nachev 2017), we excluded the first 150 nose pokes at the rewarding dispensers (Figs. S2–S5). We also tried the following alternative approaches: taking the 100 nose pokes between the 151st and the 251st, taking the last 100 or the last 20 nose pokes, or only taking the nose pokes after the discrimination performance (see below) in two consecutive blocks of 20 nose pokes exceeded the individual mean performance for that drinking session. None of the major results were qualitatively changed with the alternative cut-off points (except for experiment 3, see discussion), so in the main results we only report the results with the exclusion of the first 150 nose pokes at the rewarding dispensers.

From the remaining nose poke data we calculated the *discrimination performance* for each mouse and each condition of each experiment. Since each condition was repeated twice (initial acquisition and reversal), we calculated the discrimination performance as the total number of nose pokes at the high-profitability dispenser divided by the sum of the total number of nose pokes at the high- and at the low-profitability dispensers. Nose pokes at the non-rewarding dispensers were ignored. In the incongruent condition of experiment 2 in which the profitability was equal (relative value = 1, Fig. 2), the dispenser with the higher reward volume was treated as the “high-profitability” dispenser. It is important to emphasize that the discrimination performance does not necessarily reflect the capability of mice to distinguish between options, but also depends on other factors such as (over-)training, motivation, and exploratory behavior. Thus, the primary measure in our experiments was the discrimination performance that could develop in one drinking session, controlled for positional biases.

#### Equivalence tests in experiments 1, 2, and 4

In order to investigate how the two reward dimensions contributed towards choice in experiments 1, 2, and 4, we looked at the contrasts between the baselines (when only one dimension was relevant) to the conditions when the two dimensions were congruent or incongruent to each other. We statistically evaluated these contrasts with the two one-sided procedure (TOST) for equivalence testing (Lauzon and Caffo 2009; Lakens 2017b).

First, we picked *a priori* a smallest effect size of interest (sesoi) as the difference in discrimination performance of 0.1 units in either direction. This value was chosen based on standard deviations (sd) in discrimination performance observed in previous studies (e.g. Fig. 4 in Rivalan, Winter, and Nachev 2017), which ranged from 0.05 to 0.1. Although discrimination performance is bound by 0 and 1, most empirical values, especially the differences between two values, are far enough from these bounds so that their distribution approaches the normal. The expected sd of the difference between two normal distributions with sd of 0.1 (we conservatively picked the largest value) is 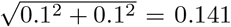. With this standard deviation and a sample size of 24, the equivalence bounds needed to detect equivalence of paired samples with a power of 0.95 are [-0.1, 0.1] (powerTOSTpaired.raw function in package TOSTER, Lakens 2017a). The sesoi can be graphically represented as the [-0.1, 0.1] interval around the difference of zero, or as [0.4, 0.6] around the chance performance of 0.5. We then estimated the mean differences and their confidence intervals (CIs) from 1000 non-parametric bootstraps using the smean.cl.boot function in package Hmisc (Harrell and Dupont 2019). For a single equivalence test the 90% CI is usually constructed, i.e. 1 − 2*α* with *α* = 0.05, because both the upper and the lower confidence bounds are tested against the sesoi (Lauzon and Caffo 2009; Lakens 2017b). This 90% CI can be fully bounded by the sesoi interval, in which case the observed effect is statistically smaller than any effect deemed worthwhile. In the opposite case there is no statistical support for equivalence. With conventional null hypothesis testing, the 95 % CI either does not include the null hypothesis (usually zero), in which case there is a statistically significant difference, or, if it does include the null, the difference is not statistically significant. When combining the equivalence and null hypothesis tests (which can also be done with examination of the 95% and 90% confidence intervals), there are four possible outcomes (Lakens 2017b):

1. If the 90% CI is fully bounded by the sesoi and the 95% CI includes the null, there is statistical support for equivalence.
2. If the 90% CI is fully bounded by the sesoi, but the 95% CI does not include the null, there is statistical support for difference with an effect size smaller than the sesoi. This result can be interpreted as practical equivalence or trivial difference.
3. If the 90% CI is not fully bounded by the sesoi, but the 95% CI includes the null, the result is deemed inconclusive.
4. If the 90% CI is not fully bounded by the sesoi and the 95% CI does not include the null, there is statistical support for difference.

Therefore, we only considered absolute differences in discrimination performance of at least 0.1 to be of practical significance in our study. Smaller differences, regardless of their statistical significance using other tests, were considered to be trivial.

#### Linear regression and equivalence tests in experiment 3

In order to test whether the background dimension affected discrimination performance we fitted linear regression models for each mouse and each dimension, with discrimination performance as the dependent variable and background level as the independent variable. The background level was the proportion of the actual value to the maximum of the four values tested, e.g. the background levels for volumes 4, 10, 15, 20 were 0.2, 0.5, 0.75, 1, respectively. We defined *a priori* a smallest effect size of interest (sesoi), as 0.125, which is the slope that would result from a difference of 0.1 (the sesoi in experiments 1, 2, and 4) in discrimination performance between the smallest and the largest background levels (PV1 and PV4, 0.2 and 1, respectively). A slope estimate (whether positive or negative) within the sesoi interval would allow us to reject an effect of background dimension of 0.125 or larger, which can be interpreted as *practically* equivalent to an absence of a meaningful effect.

#### Control of type I error rate

Researchers have shown that in order to correct for multiple comparisons in equivalence tests, it suffices to apply a familywise correction of the *α* for the problematic cases where the type I error is most likely (Davidson and Cribbie 2019), i.e. when equivalence is supported, but the mean difference is close to the sesoi bound. The families of tests, for which multiple comparisons occur in our study, are the eight contrasts in each of experiments 1, 2, and 4 (three families), the tests on the two slopes in experiment 3, and the six before-after contrasts between experiment 1 and 4. For each of these five families the *α* was divided by *k*^2^*/*4, where *k* was the number of problematic cases in each family (Caffo, Lauzon, and Röhmel 2013). However, the number of problematic cases did not exceed two in any of the test families, which resulted in the corrected *α* equal to the original value of 0.05. Furthermore, even with *k* equal to eight, two, and six (the total number of tests in each test family), only a single result changed from non-equivalent to inconclusive. We therefore report the uncorrected 90% and 95% CIs.

### Simulations

In order to examine whether the behavior of the mice was more consistent with integrative or with non-compensatory rules, we implemented simulations with six different decision rules. We based our decision models on the Scalar Utility Theory (SUT: Kacelnik and Brito e Abreu 1998; Rosenström, Wiesner, and Houston 2016), which models memory traces for reward amounts (or volumes) as normal distributions rather than point estimates. The scalar property is implemented by setting the standard deviations of these distributions to be proportional to their means. Choice between two options with different volumes can be simulated by taking a single sample from each memory trace distribution and selecting the option with the larger sample.

As previously explained, the discrimination performance for reward probabilities can be reasonably predicted by the relative intensity of the two options (Rivalan, Winter, and Nachev 2017). This suggests that the memory traces of reward probabilities also exhibit the scalar property, so that discrimination of small probabilities (e.g. 0.2 vs. 0.5, *i* = 0.86) is easier than discrimination of large probabilities (e.g. 0.5 vs. 0.8, *i* = 0.46). Consequently, discrimination (of either volumes or probabilities) when options vary along a single dimension can be predicted by SUT.

#### Virtual mice

In order to extend the basic SUT model for multidimensional choice situations, we implemented six variations that differed in the use of information from the volume and probability dimensions (Table 1), including integrative and non-compensatory models. The information from the different reward dimensions was used to obtain for each choice option a *remembered value* (utility), which exhibited the scalar property. Choice was simulated by single sampling from the *remembered value* distributions with means equal to the *remembered values* and standard deviations proportional to the *remembered values*.

**Table 1:**
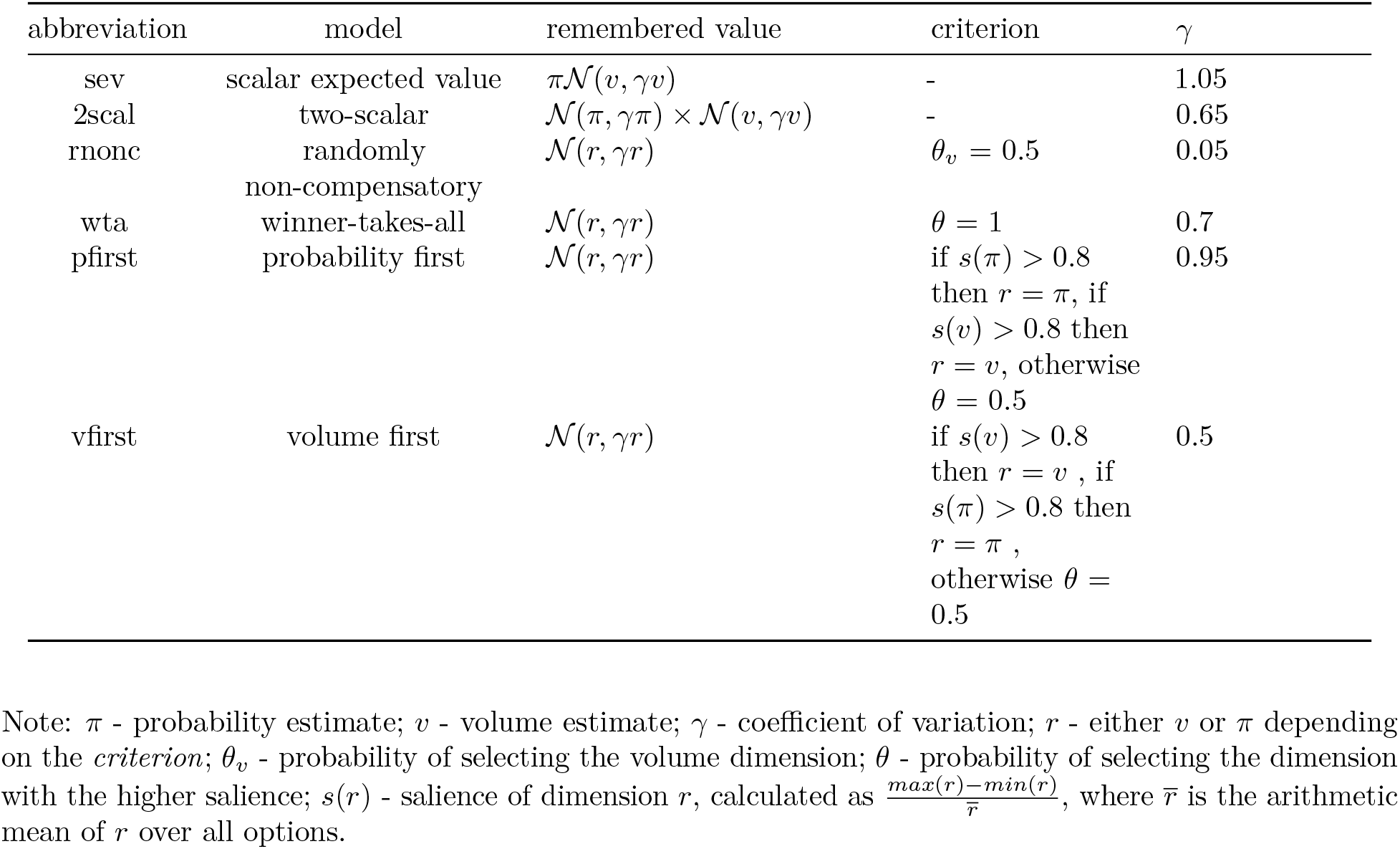
Decision-making models.

In an earlier version of the foraging model, mice started without knowledge of the reward properties and learned through Bayesian updating (Foley and Marjoram 2017). To focus on post-acquisition performance we removed the first 150 visits, like we did with the empirical data. Analyzing the *remembered values* of the virtual mice revealed that they had converged on the actual reward values with a small fluctuation around those. For simplicity, here we decided to simulate only post-acquisition discrimination performance. The virtual mice thus began each experimental condition in a learned state with *remembered values* equal to the reward dimensions for both choice options and (further) learning was not simulated. Modelling the learning process is outside the scope of this study.

From its memory traces a virtual mouse generated one *remembered value* distribution for each choice option, according to one of six different rules (see below, Table 1). Action selection was then implemented by taking a single sample from each distribution and selecting the option with the larger sample.

#### Decision-making models

##### 1. Scalar expected value model

There is a single memory trace for each option and it consists in the simple product of the estimate for the volume and the estimate for the probability (expected value). The scalar property is implemented as 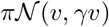, where *π* is the probability estimate. 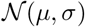 is a normal distribution with mean *μ* and standard deviation *σ*, *v* is the volume estimate, and *γ* is a free parameter, the coefficient of variation. This model thus utilizes information from all dimensions for every decision.

##### 2. Two-scalar model

There are traces for each dimension for every option, where each trace exhibits the scalar property independently and the value is obtained by simple multiplication of the traces for each dimension: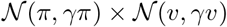. This model also utilizes information from all dimensions for every decision. Although it allows each dimension to have its own scalar factor, e.g. *γ_π_* ≠ *γ_v_*, for simplicity we assume that they are both equal.

The memory traces in the remaining models are identical to the traces in the two-scalar model, but these models usually consider only a single dimension.

##### 3. Randomly non-compensatory model

Each decision is based on a single dimension, selected with probability *θ_v_* = 0.5.

##### 4. Winner-takes-all model

Each decision is based only on the dimension with the highest salience. The salience for a vector of estimates from memory traces (mean values) along one dimension, e.g. volume *v* = (*v*_1_, *v*_2_, …, *v_n_*), is calculated as 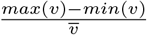, where *n* is the number of options. In the case of *n* = 2, the salience is equivalent to the previously described relative intensity measure. For dimensions of equal salience the model reverts to random choice.

The last two models are examples of a lexicographic rule, in which the dimensions are checked in a specific order. If the salience of a dimension is higher than a given threshold, then a decision is made based only on this dimension. Otherwise the next-order dimension is checked. If all dimensions have saliences below the threshold, the model reverts to random choice. The value of the threshold was set at 0.8, the psychometric function threshold for probability (Rivalan, Winter, and Nachev 2017), but we also performed sensitivity analyses on the threshold values (Fig. S8, Fig. S9).

##### 5. Probability first model

Probability is checked first, then volume.

##### 6 Volume first model

Volume is checked first, then probability.

#### Environment

Each of the experimental conditions was recreated in the simulations as a binary choice task between the high-profitability and the low-profitability options. We did not simulate the two non-rewarding options. Upon a visit by a virtual mouse, a choice option would deliver a reward with its corresponding volume and probability (Fig. 2). The virtual environment was not spatially and temporally explicit. Thus, no reversal conditions were simulated and the test of each experimental condition consisted in a sequence of 100 choices. All experimental conditions in all four experiments were tested.

#### Model fits

All models described above share the same free parameter, the scalar factor *γ*. In order to obtain baseline estimates for *γ* for each of the models (Table 1), we focused on the probability baseline discrimination performances of all mice in experiments 1 and 4 (baseline conditions BPLV and BPHV). We performed a grid search sensitivity analysis by varying *γ* with steps of 0.05 in the range of (0.05, 2). We generated 100 decisions by 100 mice for each cell in this grid and then used locally weighted scatterplot smoothing (loess) to fit a model for each condition. The free parameter values that resulted in the smallest RMSEs compared to the observed baseline data were selected for the comparison of the six models (Table 1). We also performed a sensitivity analysis for different values of the free parameters *θ_v_* in the randomly non-compensatory model and of the thresholds for volume and probability in the volume first and probability first models, in the range of (0, 1), with a step of 0.05. The resulting free parameter estimates (across animals) were then used in out-of-sample tests of the six models. For each of the experimental conditions in the four experiments (Fig. 2) and for each of the six models we simulated 100 choices by 100 (identically parametrized) mice. Over the 100 choices we calculated the discrimination performance for each mouse and then used the median of the individual discrimination performances as the model prediction. We then quantified the model fits to the empirical data by calculating root-mean-square-errors (RMSE), excluding the BPLV and BPHV conditions in experiments 1 and 4. Finally, we ranked the models by their RMSE scores.

## Results

### Experiment 1: Mice consistently preferred the more profitable option, even with a trade-off between reward probability and reward volume

Generally, compared to the baselines, mice showed an increase in discrimination performance in the congruent condition and a decrease in performance in the incongruent condition (Fig. 3a,b). The only exception was the C - BVHP contrast, which had an effect size smaller than the sesoi (0.05, 95% CI = [0.02, 0.09]). Furthermore, when we excluded cohort 2, the C - BVHP contrast became equivalent to zero (0.01, 95% CI = [-0.02, 0.04]). Contrary to our expectations based on previous work, the trade-off between volume and probability chosen for this experiment did not abolish preference for the higher volume option in the incongruent condition, with a discrimination performance significantly higher than the chance level of 0.5 (0.57, 95% CI = [0.51, 0.64], Fig. 3c). However, we again observed very different behavior in cohort 2, which showed a preference for the higher probability option (Fig. 3c). Thus, at least for mice in cohorts 1 and 3, in the incongruent condition there was a preference for the more profitable option and the subjective contrast in probability was not stronger than the subjective contrast in volume.

**Figure 3:**
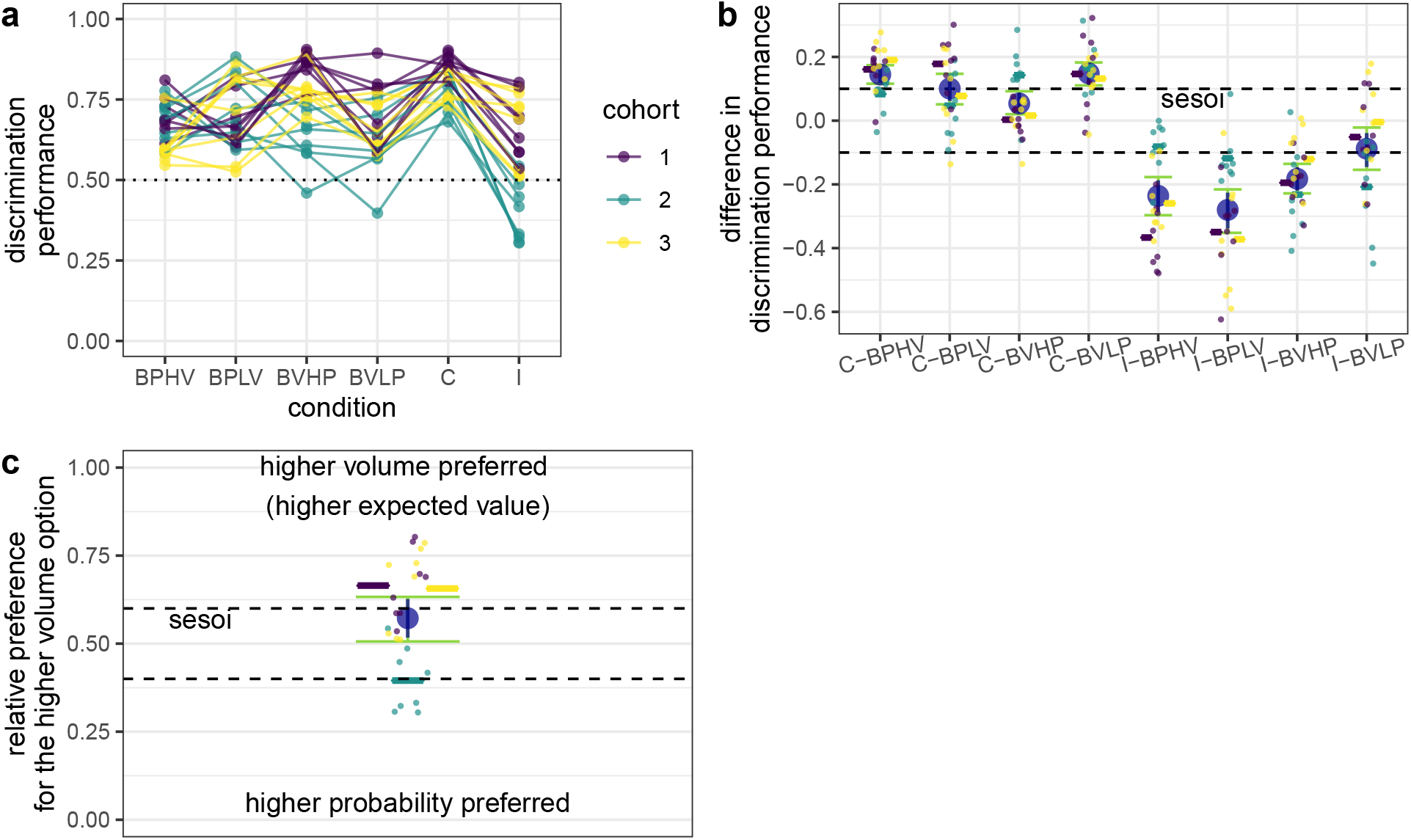
Discrimination performance in experiment 1. (**a**) Each dot is the mean discrimination performance of an individual mouse over two presentations of the same condition (initial acquisition and reversal). Experimental conditions are described in detail in Fig. 2. The discrimination performance gives the relative visitation rate of the more profitable option, or, in the incongruent condition, the option with the higher volume. Dotted line gives the chance level (0.5). Data are shown in different colors for three different cohorts of eight mice each (total *n* = 24). Data from the same individuals are connected with lines. Cohort 2 (green) was tested in a different cage set-up than the other two (see Methods for details). (**b**) Difference between discrimination performance in the baseline conditions and in the congruent and incongruent conditions. Dots show the individual differences in discrimination performance for the given conditions of each individual mouse (color-coded for cohort as in (a)). Positive differences indicate an increase in performance and negative differences - a decrease in performance, compared to the baseline. Horizontal colored lines give the cohort means. Large blue circles give the means and the blue vertical lines the 90% confidence intervals from non-parametric bootstraps. The smallest effect size of interest (sesoi) is represented by the dashed lines. Green whiskers give the 95% CI from non-parametric bootstraps. When the blue confidence intervals lie completely within the sesoi interval there is statistical support for equivalence (Lakens 2017b). The discrimination performance in the incongruent condition was calculated as the relative preference for the higher probability dispenser when contrasted with the probability baselines (e.g. I - BPLV) and for the higher volume dispenser when contrasted with the volume baselines (e.g. I - BVHP). (**c**) Discrimination performance in the incongruent condition. Dashed lines give the sesoi around chance level performance. Remaining notation is the same as in (b). In this experiment the option with the higher volume was also the more profitable option.

### Experiment 2: Some evidence for equal weighing of reward probability and reward volume

Similar to experiment 1, in experiment 2 mice showed an increase in discrimination performance in the congruent condition (with one exception) and a decrease in performance in the incongruent condition (Fig. 4a,b). This time, the exception was the C - BPLV contrast, which was equivalent to 0 (0.02, 95% CI = [-0.01, 0.04]). Although the discrimination performance in the incongruent condition was again different from 0.5 (0.41, 95% CI = [0.35, 0.47]), it was lower than chance, thus skewed towards probability (Fig. 4b). However, when the data from cohort 2 were excluded, the discrimination performance became equivalent to 0.5 (0.48, 95% CI = [0.42, 0.54]). We return to the differences between cohorts in the discussion. Thus, it appears that, at least for mice in cohorts 1 and 3, the subjective contrasts in volume and probability were equal and no reward dimension seemed to have priority over the other.

**Figure 4:**
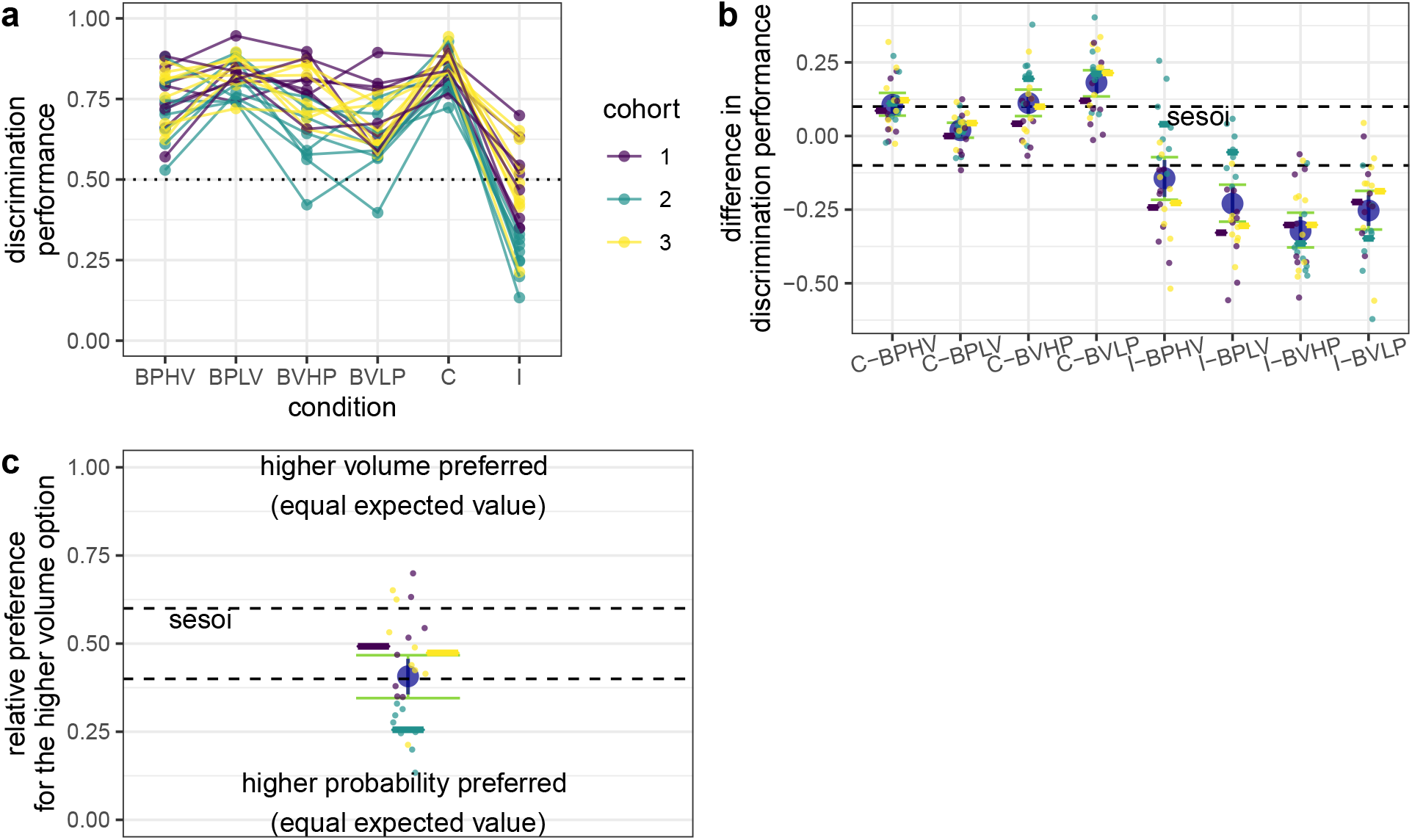
Discrimination performance in experiment 2. Same notation as in Fig. 3. (**a**) Discrimination performance in all conditions. (**b**) Difference between discrimination performance in the baseline conditions and in the congruent and incongruent conditions. The discrimination performance in the incongruent condition was calculated as the relative preference for the higher probability dispenser when contrasted with the probability baselines (e.g. I - BPLV) and for the higher volume dispenser when contrasted with the volume baselines (e.g. I - BVHP). (**c**) Discrimination performance in the incongruent condition. In this experiment both options were equally profitable and had the same expected value.

### Experiment 3: Probability discrimination decreased with an increase in reward volume, but volume discrimination was not affected by changes in reward probability

The results of experiment 3 show that the discrimination performance for probability decreased with increasing volumes, although the effect size was small (−0.1, 95% CI = [−0.16, −0.06], without cohort 2: (−0.147, 95% CI = [−0.212, −0.088], Fig. 5). In contrast, the discrimination performance for volume was *practically* independent from probability as the background dimension, since the estimate for the slope was smaller than the sesoi (0.06, 95% CI = [−0.01, 0.14], Fig. 5). Without cohort 2 the slope estimate for the volume dimension was still small, but significantly positive (0.072, 95% CI = [0.009, 0.139]). These results partially support the hypothesis that decision-makers may ignore a reward dimension along which options do not vary.

**Figure 5:**
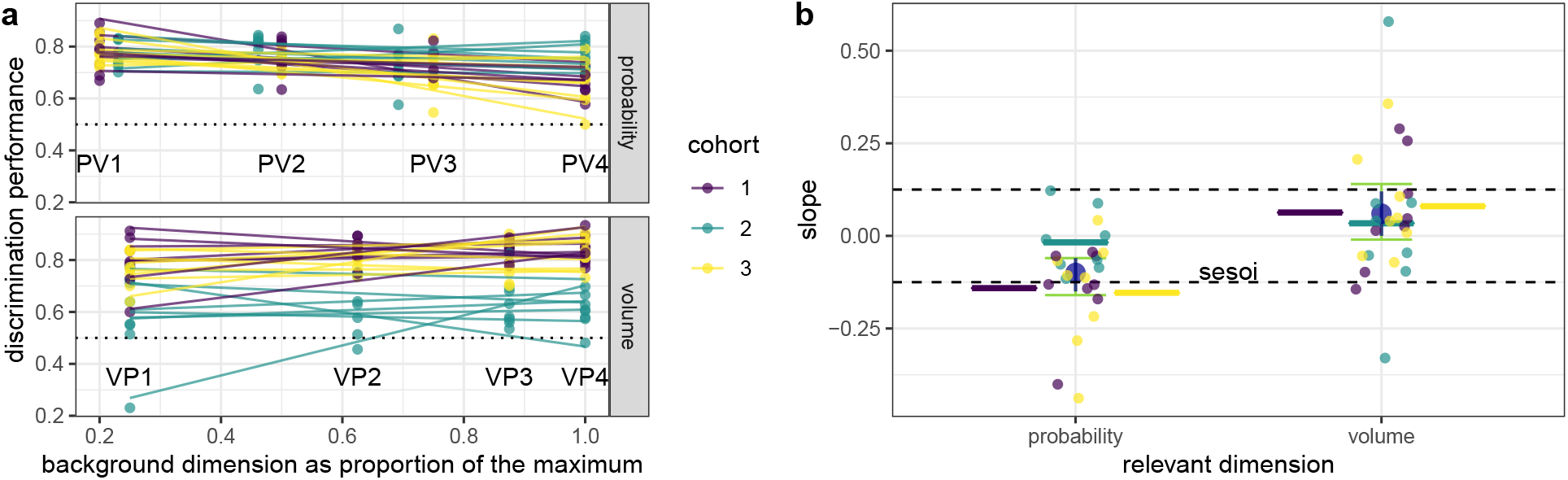
Effect of background dimension on discrimination performance in experiment 3. (**a**) The two choice options always differed along the relevant dimension either probability or volume (panels). The discrimination performance for each mouse was measured at four different levels of the background dimension, which was set at the same values on both rewarding options during a single drinking session, but differed from condition to condition (Fig. 2). Each dot (color-coded for cohort number) is the mean discrimination performance of an individual mouse over two presentations of the same condition (initial acquisition and reversal). Dotted line gives the chance level of 0.5. Data are shown in different colors for three different cohorts of eight mice each (total *n* = 24). Lines give best linear fits. Cohort 2 (green) was tested in a different cage set-up than cohorts 1 and 3 (see Methods for details). (**b**) Each colored dot represents the individual slope of one line in (a). The smallest effect size of interest (sesoi, dashed lines) was determined to be the slope (0.125) that would have resulted in a difference in discrimination performance of 0.1, from the lowest to the highest level of the background dimension (from PV1 to PV4 in (a)). Large blue circles give the means and the blue vertical lines the 90%-confidence intervals from non-parametric bootstraps. Green whiskers give the 95% CI from non-parametric bootstraps. Horizontal colored lines give the cohort means.

### Experiment 4: Mice improved their volume discrimination over time

In the comparison between experiment 1 and experiment 4, mice showed an improved discrimination performance in both volume baselines, as well as in the incongruent and BPLV conditions (Fig. 6). There was only a trivial improvement in the congruent condition (Fig. 6). When we applied a familywise error control procedure, only the BPLV result changed from an increase to inconclusive and the congruent condition, from trivial to equivalent. Thus, consistent with our prior findings, mice improved their volume discrimination over time.

**Figure 6:**
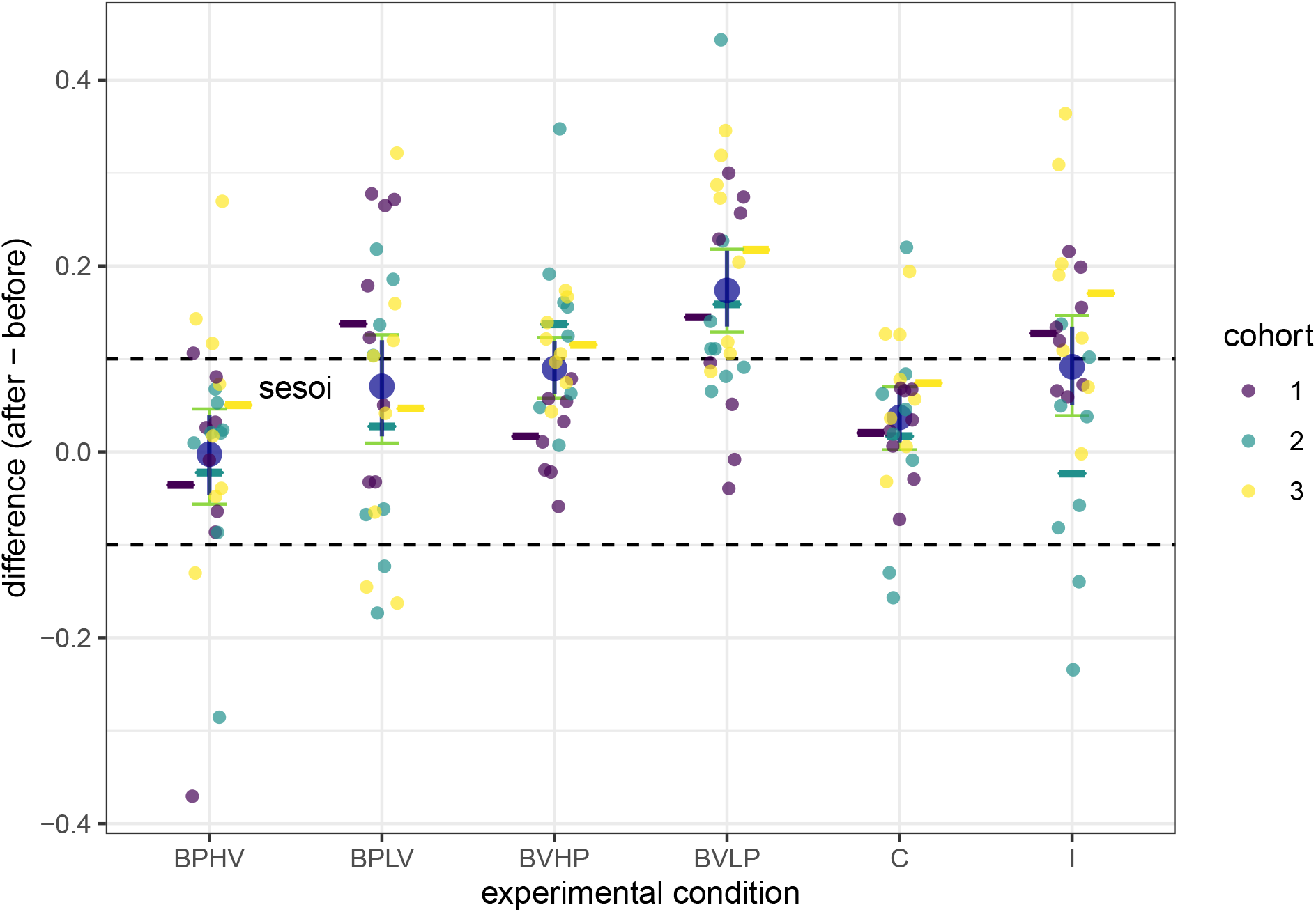
Difference in discrimination performance between identical conditions in experiment 1 and experiment 4. Same notation as in in Fig. 3. The sequence of conditions was pseudo-random in each experiment and different for each individual. Positive differences indicate an increase in discrimination performance with time. Mice were seven weeks old at the beginning of experiment 1 and 13-14 weeks old at the beginning of experiment 4. The discrimination performance in the incongruent condition was calculated as the relative preference for the higher volume dispenser.

The discrimination performance in the congruent condition was better than either of the probability baselines, but equivalent to the volume baselines (Fig. 7a,b). For cohorts 1 and 3, the discrimination performance in the incongruent condition was lower than in any of the four baselines, but the difference from the volume baselines was smaller (Fig. 7b). Cohort 2 showed the opposite pattern (Fig. 7b). Finally, compared to experiment 1 the influence of the volume dimension on choice and the discrepancy between cohort 2 and the other cohorts were even more pronounced (Fig. 7c).

**Figure 7:**
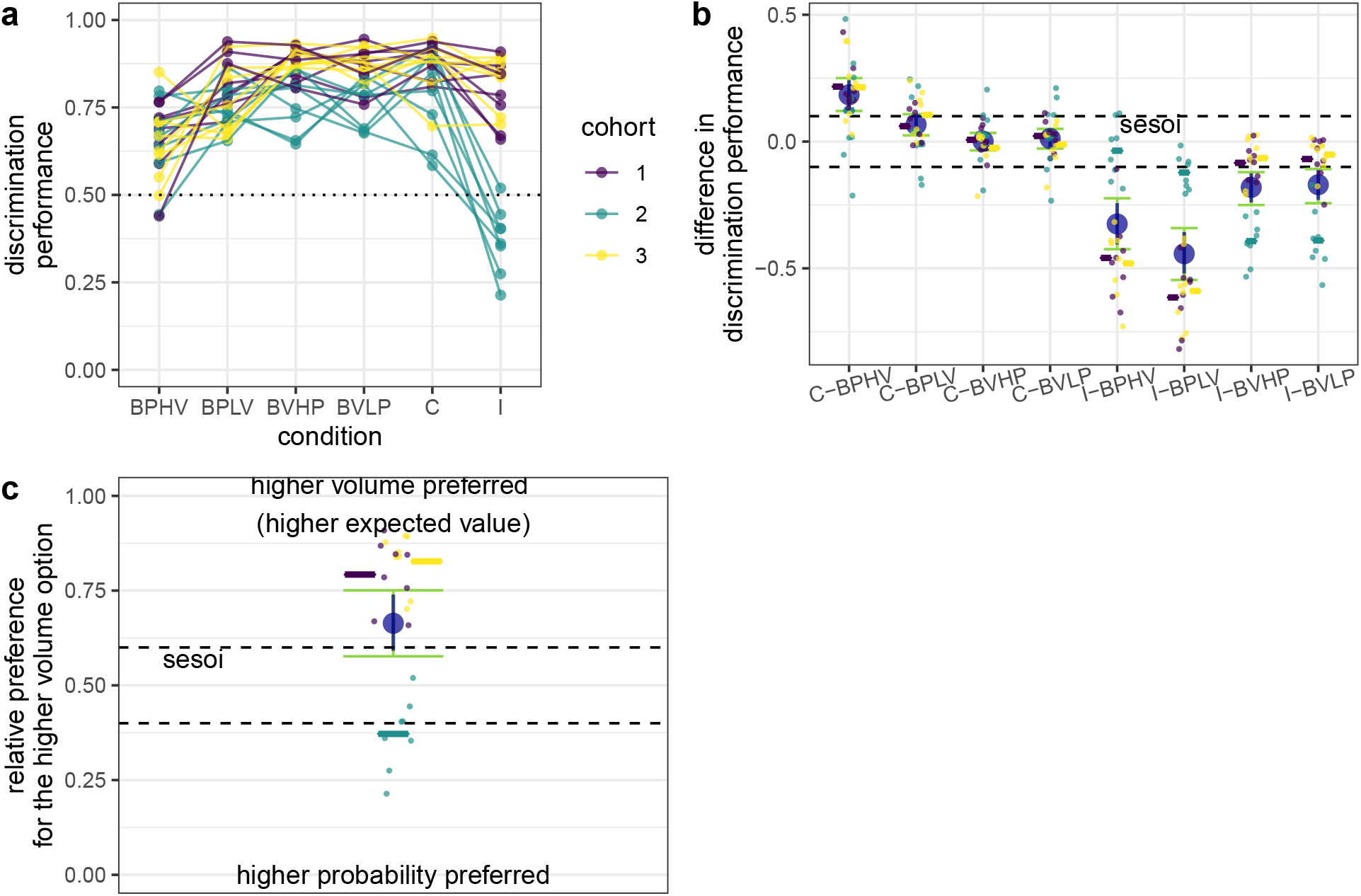
Discrimination performance in experiment 4, with identical conditions to experiment. Same notation as in Fig. 3. (**a**) Discrimination performance in all conditions. (**b**) Difference between discrimination performance in the baseline conditions and in the congruent and incongruent conditions. The discrimination performance in the incongruent condition was calculated as the relative preference for the higher probability dispenser when contrasted with the probability baselines (e.g. I - BPLV) and for the higher volume dispenser when contrasted with the volume baselines (e.g. I - BVHP). (**c**) Discrimination performance in the incongruent condition. In experiments 1 and 4 the option with the higher volume was also the more profitable option. Compare to Fig. 3.

### Decision models of two-dimensional choice suggest that mice initially relied on both reward volume and reward probability, but then developed a bias for reward volume

There was no single model that could best explain the choice of the mice in all four experiments, but the scalar expected value, two-scalar, and winner-takes-all models were in the top-three performing models most frequently (Tables 1, 2, Figs. S10–S13). However, due to the unexpected differences in performance between cohort 2 and the other cohorts (e.g. Fig. S13), we also ranked the models separately for the different mouse groups, depending on which cage they performed the experiments in (cohorts 1 and 3 in cage 1 and cohort 2 in cage 2). Indeed, two different patterns emerged for the different cages. For the two cohorts in cage 1, scalar expected value and two-scalar were the best supported models, followed by the winner-takes-all and volume first models (Table 3). Notably, the volume first model was the best performing model in the later experiments 3 and 4, but the worst model in the earlier experiments 1 and 2. In contrast, the probability first model was the best supported model for cohort 2, followed by the randomly non-compensatory, winner-takes-all, and scalar expected value models (Table 4).

**Table 2:** Best performing models ranked by root-mean-square-errors (RMSE).

| experiment |  |  |  |  |
| --- | --- | --- | --- | --- |
| rank | 1 | 2 | 3 | 4 |
| 1 | sev | sev | vfirst | 2scal |
| 2 | 2scal | 2scal | sev | wta |
| 3 | wta | wta | 2scal | sev |
| 4 | rnonc | pfirst | wta | vfirst |
| 5 | pfirst | rnonc | pfirst | rnonc |
| 6 | vfirst | vfirst | rnonc | pfirst |

**Table 3:** Best performing models ranked by root-mean-square-errors (RMSE) for cohorts 1 and 3.

| experiment |  |  |  |  |
| --- | --- | --- | --- | --- |
| rank | 1 | 2 | 3 | 4 |
| 1 | sev | 2scal | vfirst | vfirst |
| 2 | 2scal | sev | sev | 2scal |
| 3 | wta | wta | 2scal | wta |
| 4 | rnonc | rnonc | wta | sev |
| 5 | pfirst | pfirst | rnonc | rnonc |
| 6 | vfirst | vfirst | pfirst | pfirst |

**Table 4:** Best performing models ranked by root-mean-square-errors (RMSE) for cohort 2.

| experiment |  |  |  |  |
| --- | --- | --- | --- | --- |
| rank | 1 | 2 | 3 | 4 |
| 1 | pfirst | pfirst | pfirst | pfirst |
| 2 | rnonc | rnonc | wta | rnonc |
| 3 | sev | sev | 2scal | wta |
| 4 | wta | 2scal | sev | 2scal |
| 5 | 2scal | wta | rnonc | sev |
| 6 | vfirst | vfirst | vfirst | vfirst |

## Discussion

The foraging choices of the mice in this study provide evidence both for and against full integration of reward volume and probability. In the first two experiments, most mice differed in discrimination performance (increased or decreased) in the conditions in which both reward dimensions were simultaneously relevant (congruent and incongruent conditions) compared to the baselines, in which only one of the two dimensions was relevant at a time (Figs. 3, 4). Consequently, the best supported models for these two experiments (cohort 2 excluded, see discussion about differences between cohorts below) were the models that made use of the full information from both reward dimensions (sev, 2scal), or from the dimension that was subjectively more salient (wta, Table. 3). Although these models were good predictors of choices in experiments 3 and 4 as well, the best-performing model in experiments 3 and 4 was the one that considered the volume dimension first and the probability dimension only if differences on the volume dimension were insufficient to reach a decision (Table 3). Thus, it appears that mice initially used information from all reward dimensions without bias and with experience started to rely more on one reward dimension and disregarded the other when both dimensions differed between choice options. Interestingly, in human development the use of integrative decision rules has also been shown to decrease with age (Jansen, Duijvenvoorde, and Huizenga 2012).

In similar and more complex choice situations when options vary on several dimensions, an animal has no immediate method of distinguishing the relevant from the background dimensions. Instead it must rely on its experience over many visits before it can obtain information about the long-term profitability associated with the different reward dimensions. Under such circumstances a decision rule that considers all or the most salient reward dimensions initially and prioritizes dimensions based on gathered experience can be profitable without being too computationally demanding. Indeed, with the particular experimental design in this study, a mouse using a “volume first” priority heuristic would have preferentially visited the more profitable option (whenever there was one) in every single experimental condition, including the incongruent conditions.

### Scalar property considerations

An alternative explanation of our main results is that the mice used the “volume first” heuristic from the beginning of the experiment, but only became better at discriminating volumes (their coefficient of variation *γ* decreased) in the last two experiments. This interpretation is supported by the comparison between experiments 1 and 4 (Fig. 6), as well as from previous experiments (Rivalan, Winter, and Nachev 2017), in which mice improved their volume discrimination over time. However, it is not possible with these data to distinguish whether the effect was caused by training or maturation. Perhaps an increase in mouth capacity (Vora, Camci, and Cox 2016) or, potentially, in the number of acid-sensing taste receptors (Zocchi, Wennemuth, and Oka 2017) due to growth and maturation could allow adult mice to better discriminate water volumes. We assumed that mice consumed all water without spilling, but perhaps less experienced mice spill some water. Alternatively, with prolonged training mice might transition from goal directed strategies to egocentric or habitual responses (Packard and McGaugh 1996; Kosaki, Pearce, and McGregor 2018; and in mice: Kleinknecht et al. 2012). Comparing the discrimination performance of older untrained and younger trained mice would help clarify this confound.

The increase in discrimination performance for volume between experiments 1 and 4 (Fig. 6) suggests that the scalar property only approximately holds, and that the *γ* (coefficient of variation) for volume is not truly constant over a long period of time. This can be seen as evidence against the scalar expected value model, which assumes that the same coefficient of variation affects performance along each reward dimension. Instead, the improving volume discrimination supports a version of the two-scalar model, in which there are two different scalars (*γ_π_ γ_v_*). Alternatively, there might be only one scalar, associated with dynamic relative weights of the two dimensions (which can be implemented as a changing *θ_v_* in the randomly non-compensatory model, Fig. S7). Yet another model extension that can account for the improving volume discrimination would be to introduce an explicit sampling (exploration-exploitation balance) method (Sih and Del Giudice 2012; Nachev and Winter 2019). In natural conditions reward dimensions rarely remain stable over time and foragers can benefit from making sampling choices to gather information about the current state of the environment. Thus, not all choices need to be based on expected values and individuals may differ in their sampling rates (Sih and Del Giudice 2012; Rivalan, Winter, and Nachev 2017; Nachev and Winter 2019). With such an implementation it is not the scalar but the frequency of sampling visits that changes over time, causing differences in discrimination performance. The biggest challenge is that when it comes to volumes and probabilities, no direct method of interrogating an animal’s estimate and coefficient of variation exist, so that researchers have to infer these values from choice behavior, which is also affected by motivation, learning, and sampling frequency. In contrast, when it comes to time intervals, the peak procedure gives us a more direct measurement of the time estimation of animal subjects (Kacelnik and Brito e Abreu 1998).

### Interaction between dimensions and non-compensatory decision-making

Although mice were practically equally good at discriminating volume rewards at each different probability, the discrimination of probabilities decreased at higher volumes (Fig. 5; the estimated effect size was a decrease of 0.12 between a volume background at 4 *μL* and at 20 *μL*). This suggests that the two dimensions interact with each other. Absolute reward evaluation (Shafir 1994; Shafir and Yehonatan 2014) and state-dependent evaluation (Schuck-Paim, Pompilio, and Kacelnik 2004) are both consistent with this decrease in discrimination performance, but not with the small positive effect in the conditions in which the probability was the background dimension. With comparable expected values (Fig. 2) between the two series of conditions, these hypotheses make the same predictions regardless of which dimension is relevant and which is background. An alternative explanation is that arriving at a good estimate of probability requires a larger number of visits and when the rewards are richer (of higher volume), mice satiate earlier and make a smaller total number of visits, resulting in poorer estimates of the probabilities and poorer discrimination performance. Consistent with this explanation, mice made on average (± SD) 474 ± 199 nose pokes at the relevant dispensers at 4 *μL*, but only 306 ± 64 nose pokes at 20 *μL* (Figs. S1,S4: PV1 and PV4, respectively). Furthermore, when we controlled for the number of nose pokes by only analyzing the nose pokes between the 151st and 251st, the effect of volume on probability discrimination became equivalent to zero, suggesting that further learning after the 150st nose poke could have led to an improved discrimination performance. At the same time, controlling for the number of nose pokes also led to a significant (but small) positive effect of probability on volume discrimination (slope estimate 0.9, 95%CI = [0.01, 0.18]). This also suggests that at 0.2 probability it took mice more than 150 nose pokes to reach the same discrimination performance for volumes observed at probabilities higher than 0.2 (Fig. S4). These were the only qualitative changes caused by taking an alternative cut-off point rather than simply removing the first 150 nose pokes to the rewarding dispensers. As mentioned earlier, researchers have proposed that with absolute reward evaluation the difference/mean ratio in an experimental series like our experiment 3 should decrease with the increase of the background dimension, leading to a decrease in the proportional preference for the high-profitability alternative, i.e. discrimination performance (Shafir and Yehonatan 2014). However, this is only the case if the difference is calculated from the relevant dimension and divided by the mean utility. We suggest that both the difference and the mean should be calculated from the same entity, either utility or one of the reward dimensions. When, as in our sev and 2scal models 1, we calculate utility by multiplying the estimates for each dimension together, the difference/mean ratio of the utility does not change with the change in the background dimension between treatments. In fact, none of our models in experiment 3 exhibited an effect of the background dimension on the discrimination performance, with all slopes equivalent to zero (Fig. S14). Thus, our results also show that absolute reward evaluation does not necessarily predict an effect of background dimension on discrimination performance.

### Difference between cohorts

Our results revealed some striking differences in behavior between cohort 2 and cohorts 1 and 3 (most obvious in Fig. 7). The most likely explanation for this is an effect of the specific experimental apparatus. As explained in Methods, the precision of the reward volumes was lower in cage 2, which housed cohort 2. However, it is unlikely that such a small magnitude of the difference (0.33 ± 0.03 *μLstep^−^*^1^ in cage 1 vs. 1.56 ± 0.24 *μLstep^−^*^1^ in cage 2) could influence volume discrimination to the observed extent. Future experiments can address this issue by specifically manipulating the reliability of the volume dimension using the higher-precision pump. Instead, we suspect that the difference between cohorts might have been caused by the acoustic noise and vibrations produced by the stepping motors of the pumps. The pump in cage 1 was much louder, whereas the one in cage 2 was barely audible (to a human experimenter). This could have made it harder for mice in cage 2 to discern whether a reward was forthcoming, which could have influenced their choices (Ojeda, Murphy, and Kacelnik 2018). As a result, mice in cage 2 waited longer before leaving the dispenser during unrewarded nose pokes (Fig. S15). This potentially costly delay might have increased the relative importance of the probability dimension (decreased *θ_v_*), resulting in the observed discrimination performance in cohort 2. Furthermore, the same line of reasoning can also explain the improving volume discrimination: from the first to the fourth experiment there was a shift towards shorter unrewarded nose poke durations in the loud cage (cohorts 1 and 3, Fig. S15), suggesting that mice had learned over time to abort the unrewarded visits. This could have decreased the relative importance of the probability dimension (increased *θ_v_*), resulting in better volume discrimination. In an unrelated experiment we tested two cohorts of mice in both cages simultaneously and then translocated them to the other cage. The results demonstrated that differences in discrimination performance were primarily influenced by cage and not by cohort (Nachev, in prep.). Thus, the sound cue associated with reward delivery may be an important confounding factor in probability discrimination in mice, as it provides a signal for the reward outcome (Ojeda, Murphy, and Kacelnik 2018).

### Conclusion

In summary, our results show that mice could integrate reward volume and reward probability, which allowed them to select the more profitable option when the two reward dimensions varied independently. The resulting partial preference was consistent with Scalar Utility Theory. However, we also found that, with time mice improved their performance in volume (but not as much in probability) discrimination tasks and their choices became more consistent with a non-compensatory decision rule, in which volume is evaluated before probability. Finally, we found that mice could discriminate the same pair of probabilities better when reward volumes were smaller, but changes in the reward probability did not seem to affect their volume discrimination performance.

## Electronic Supplementary Material

**Figure S1:**
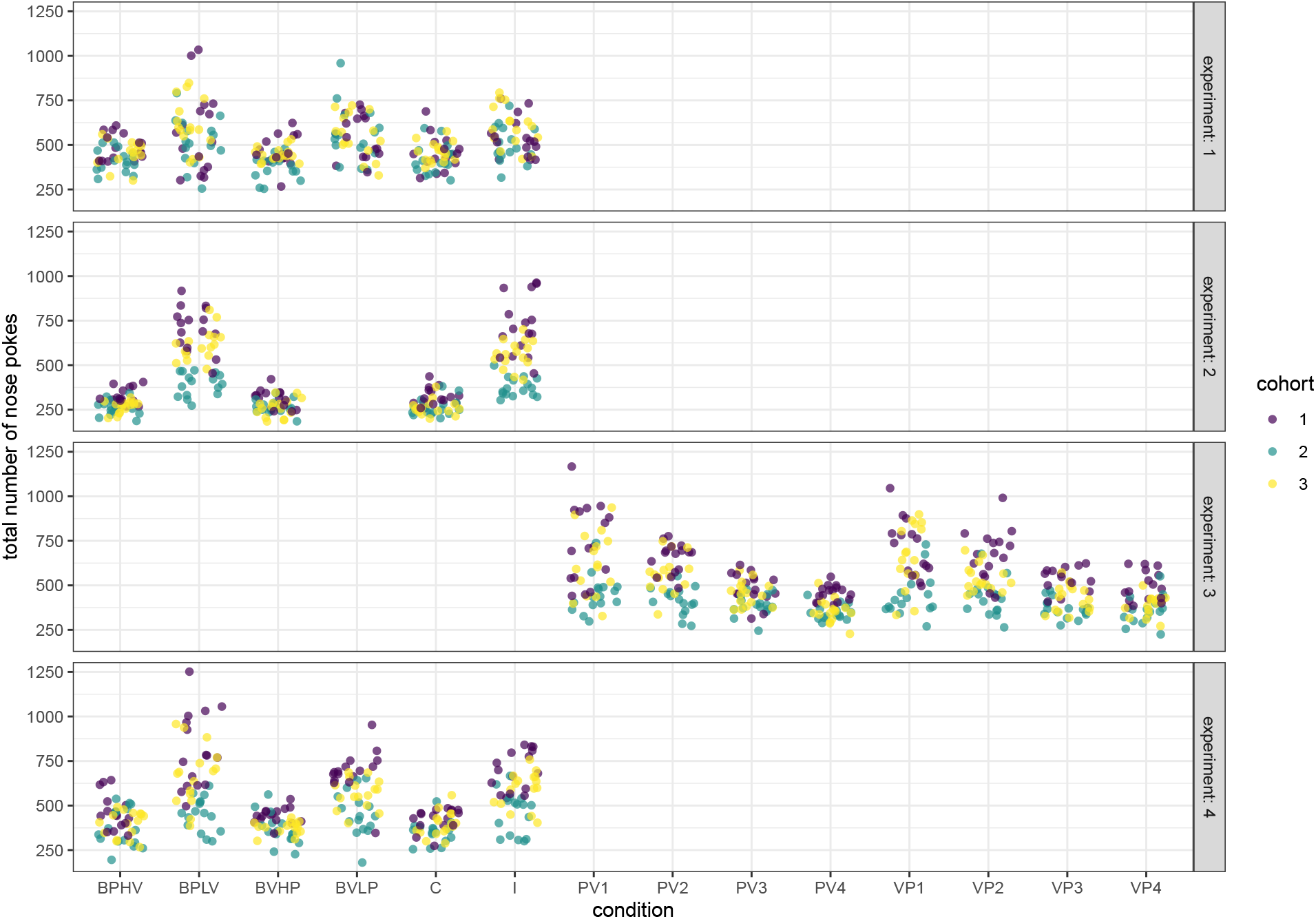
Total number of nose pokes for each experimental condition in the three cohorts in all experiments. Rows show different experiments (1-4). Each symbol represents the total number of nose pokes for a single mouse over one of the two experimental days of the given condition.

**Figure S2:**
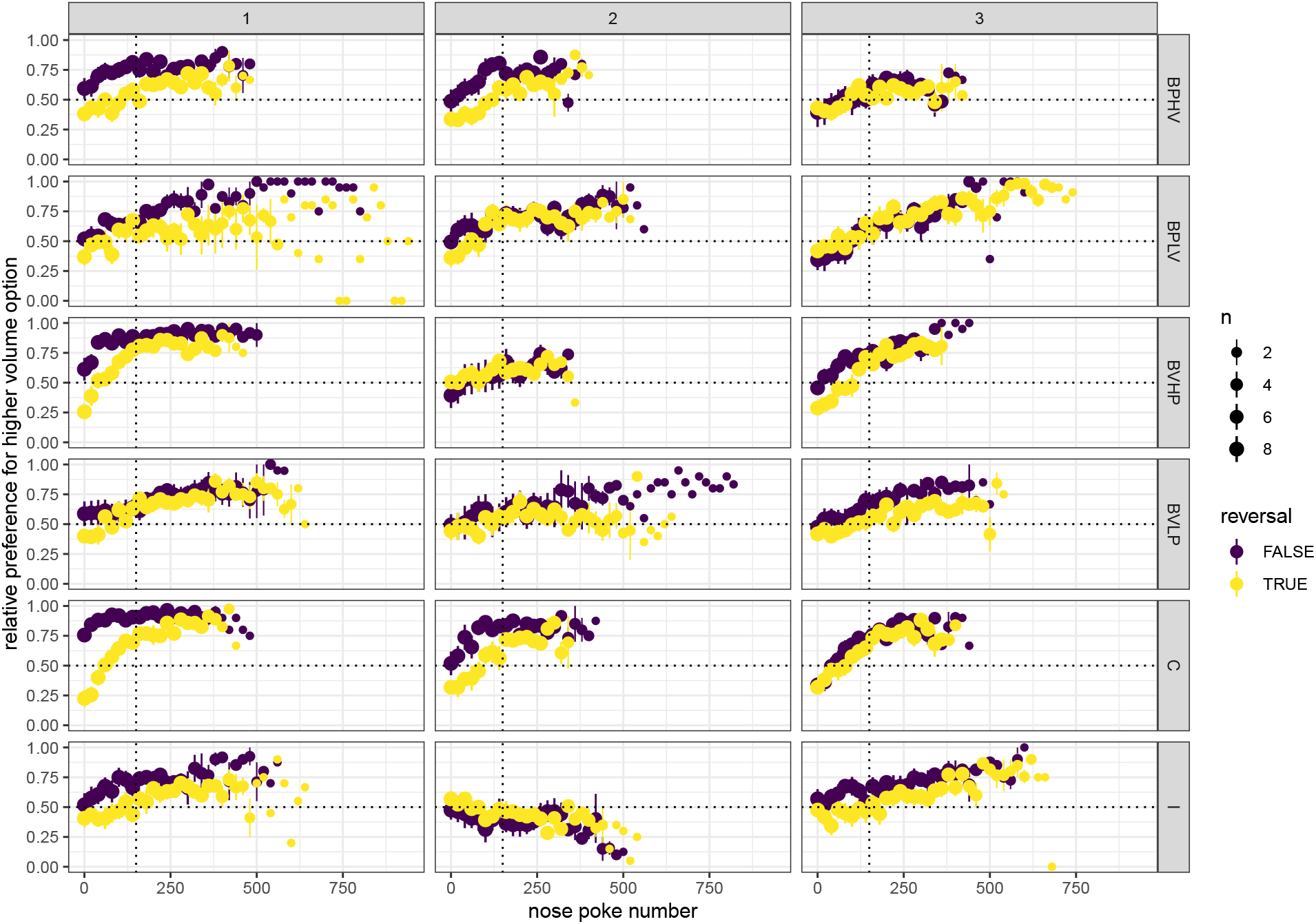
Learning curves in experiment 1. Only nose pokes at the rewarding dispensers were included (abscissa). Dots give the mean discrimination performance for blocks of 20 nose pokes over the respective number of mice (dot size) and the error bars give the standard errors. Purple symbols correspond to the first acquisition of a new condition (rows) and yellow symbols correspond to the reversal day of the same condition. The columns give the different cohort numbers (1-3). The horizontal dotted line corresponds to chance performance (0.5). The vertical dotted line corresponds to the data exclusion criterion used in the main analyses (150 nose pokes to the rewarding dispensers). For the main analyses only data to the right of this line were analysed and the purple and yellow data were pooled for each mouse, in order to calculate the discrimination performance for each condition. In most conditions the discrimination performance from the initial acquisition and reversal converges to similar values, indicating that the mice were sensitive to the reward properties and not only the location of the dispensers.

**Figure S3:**
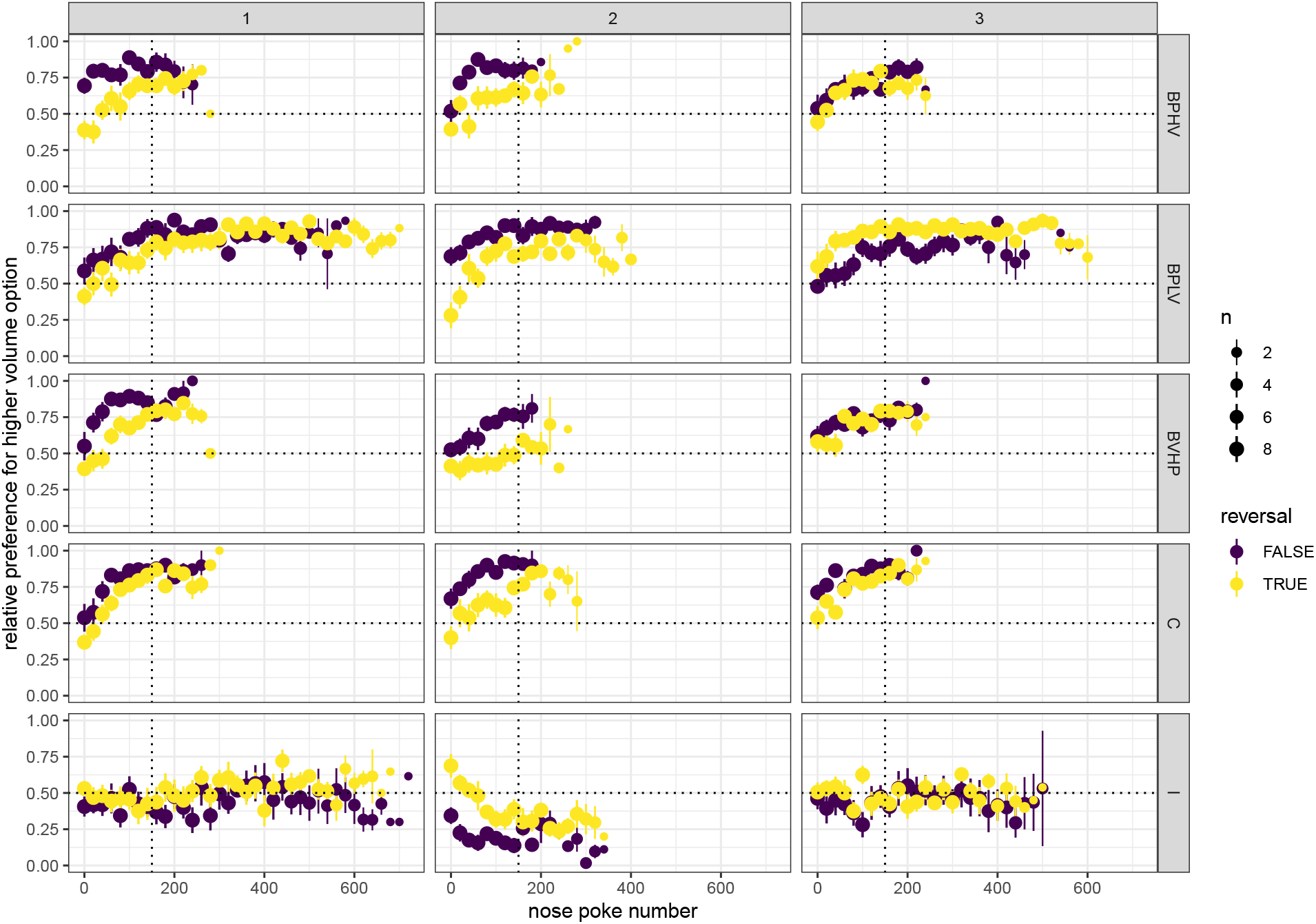
Learning curves in experiment 2. Same notation as in Fig. S2.

**Figure S4:**
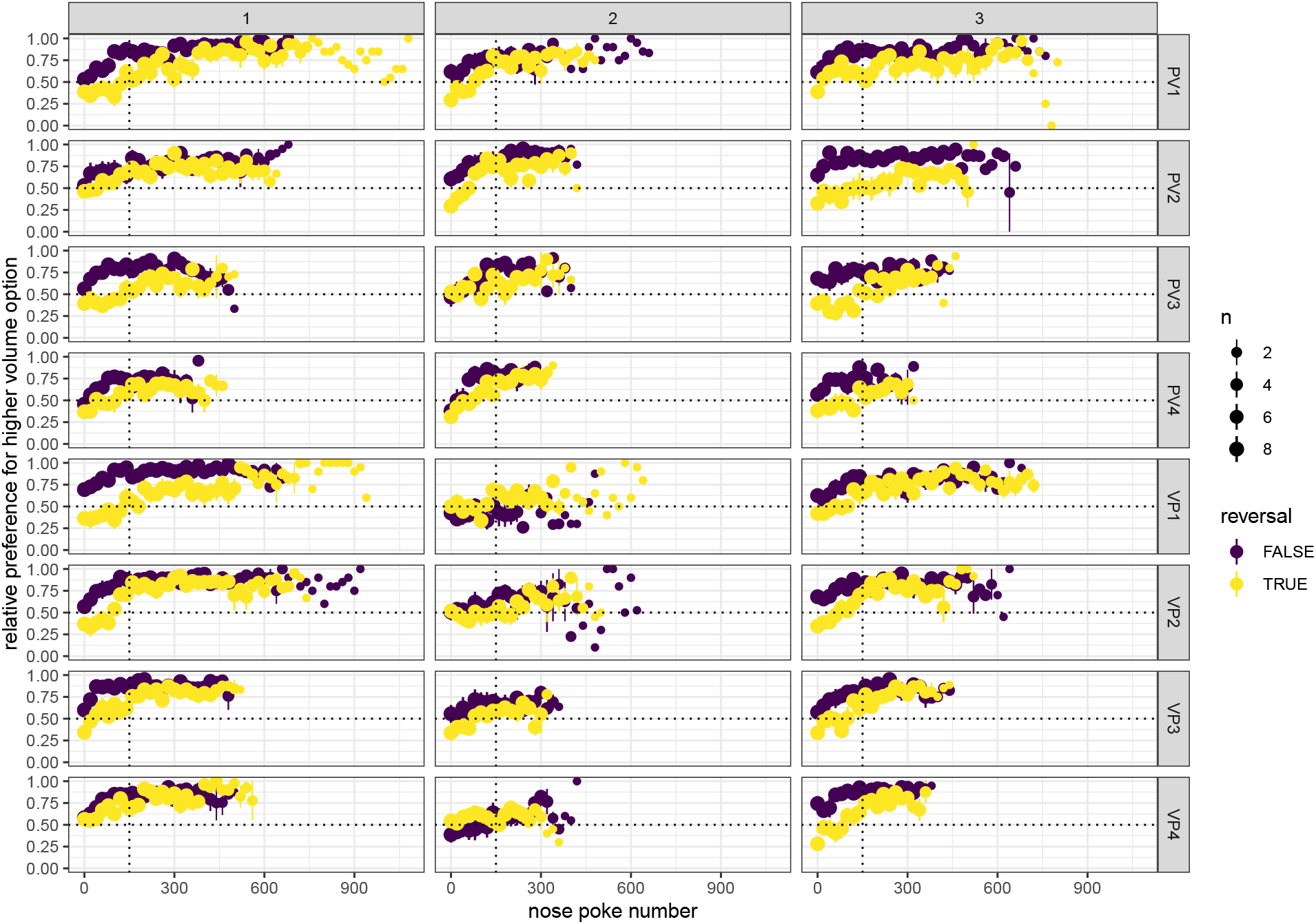
Learning curves in experiment 3. Same notation as in Fig. S2.

**Figure S5:**
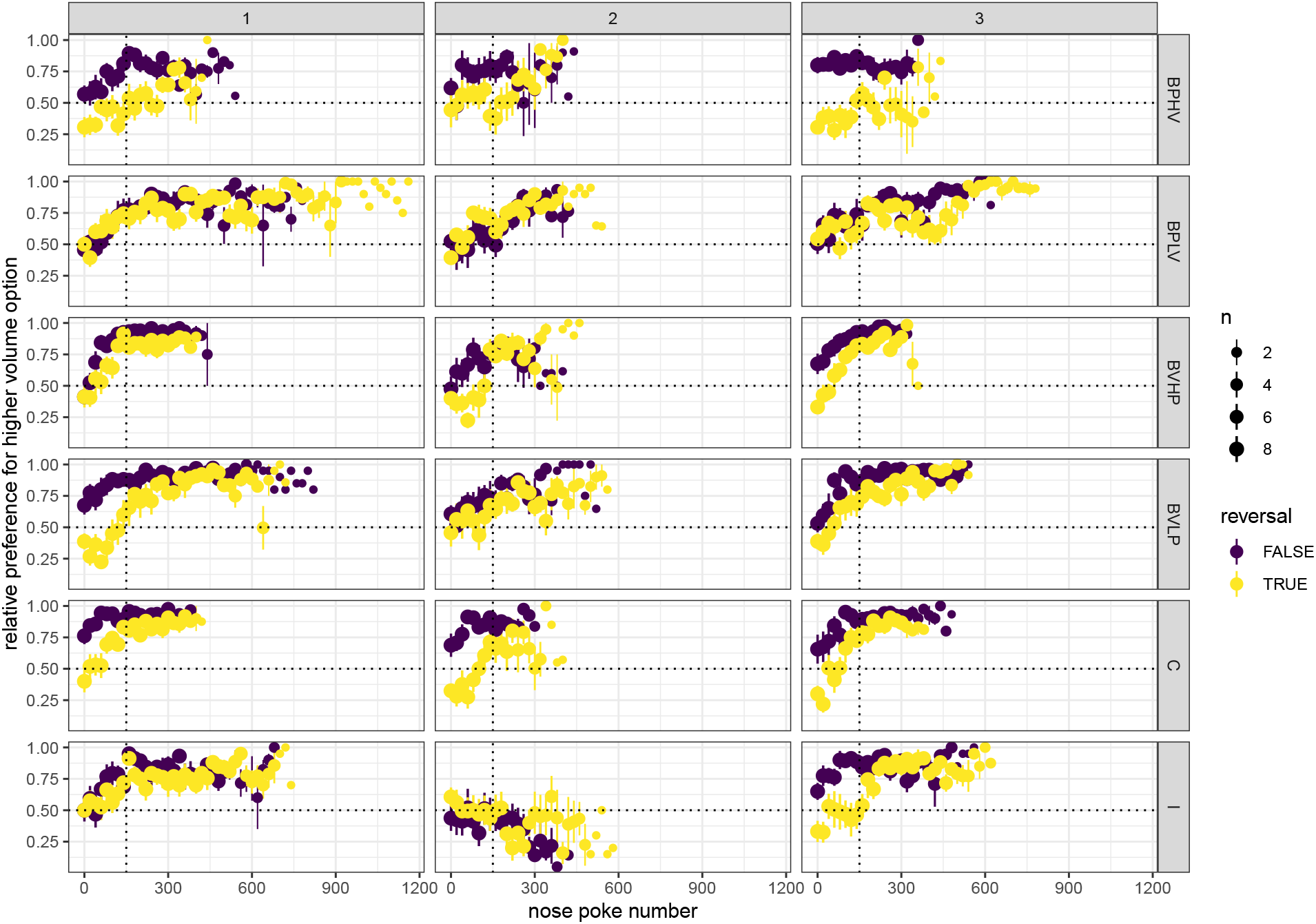
Learning curves in experiment 4. Same notation as in Fig. S2.

**Figure S6:**
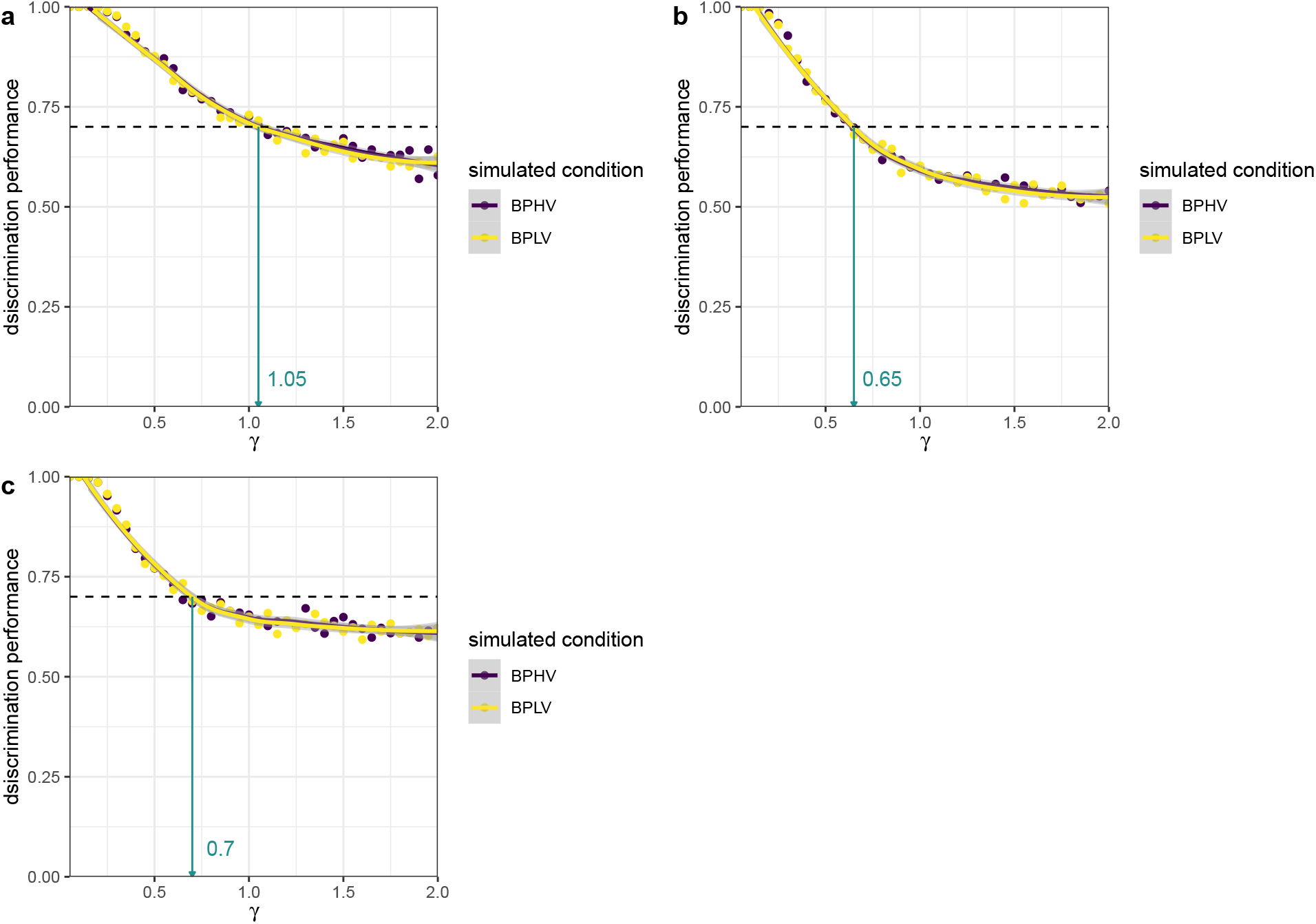
Sensitivity tests for the models that only had *γ* as a free parameter. Dots give the discrimination performances calculated from 1000 choices for each value of *γ* tested [0.05, 2] and for each of the baseline conditions BPLV (purple) and BPHV (yellow). Lines give the corresponding fits based on locally weighted scatterplot smoothing (loess). The dashed line gives the empirical mean discrimination performance from the baseline conditions BPLV and BPHV and the green arrows point to the value of gamma that resulted in the smallest root-mean-square-errors (RMSEs). These values were then used in the main simulations (Table 1). The different panels give the results for the scalar expected value (**a**), two-scalar (**b**), and winner-takes-all (**c**) models.

**Figure S7:**
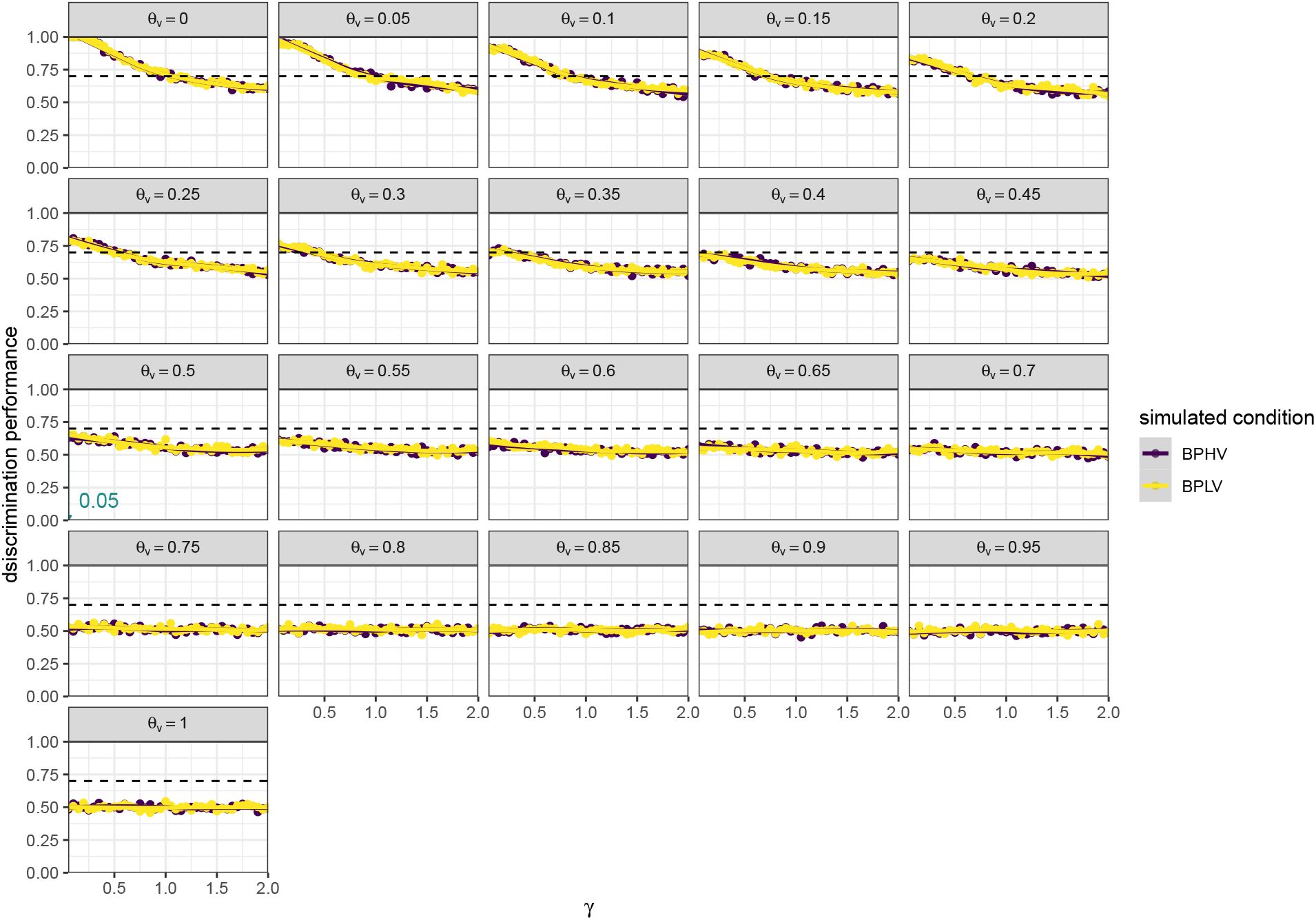
Sensitivity tests for the randomly non-compensatory model. Same notation as in Fig. S6. The different panels give the different values of the probability with which the volume dimension was chosen (*θ_v_*). For a non-biased randomly non-compensatory model we set *θ_v_* = 0.5.

**Figure S8:**
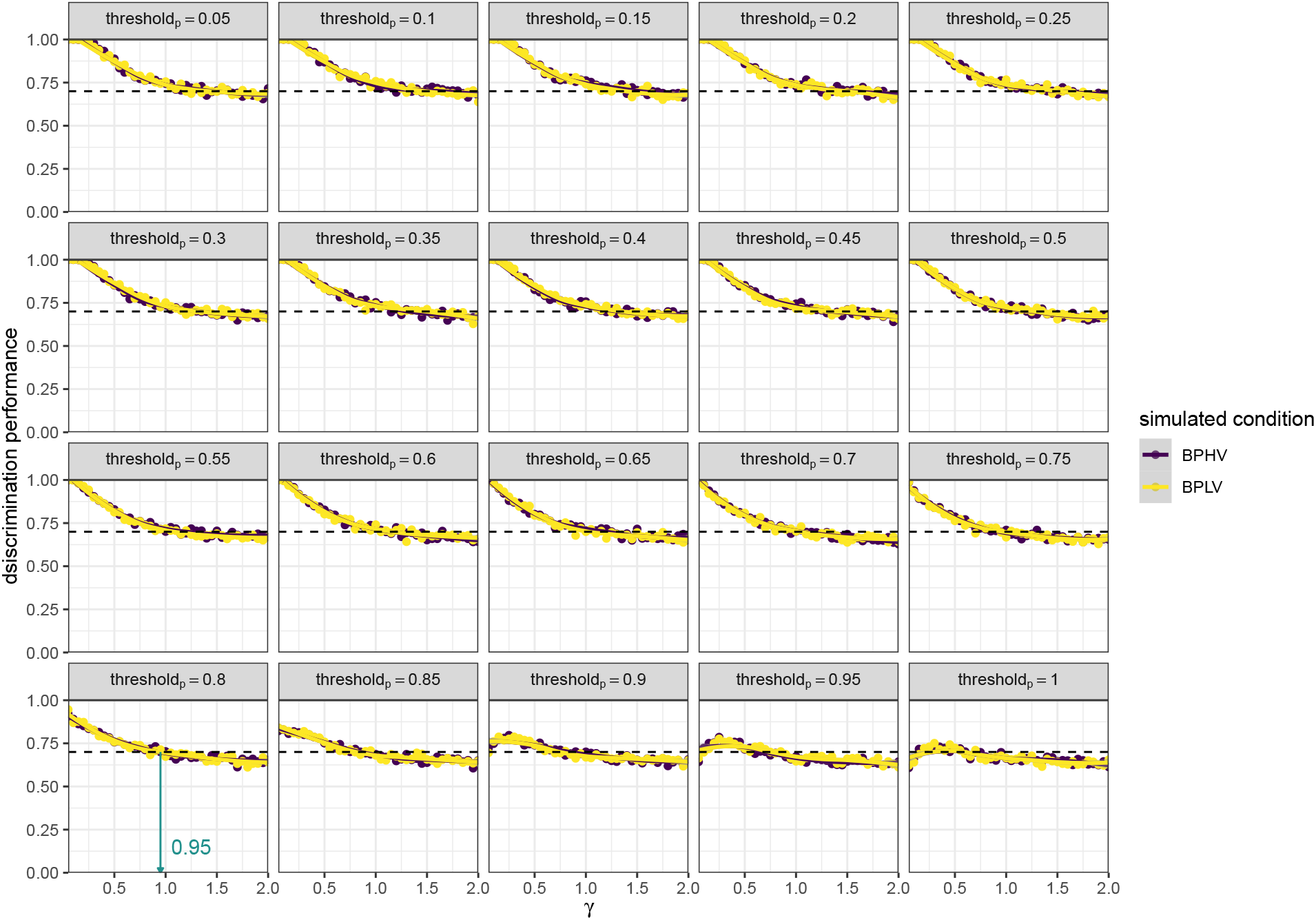
Sensitivity tests for the probability first model. Same notation as in Fig. S6. The different panels give the different values of the salience threshold that needed to be reached for one option to be preferred over the other. We set the value of the threshold for both the volume and probability dimensions to 0.8, based on the psychometric function threshold for probability (Rivalan, Winter, and Nachev 2017).

**Figure S9:**
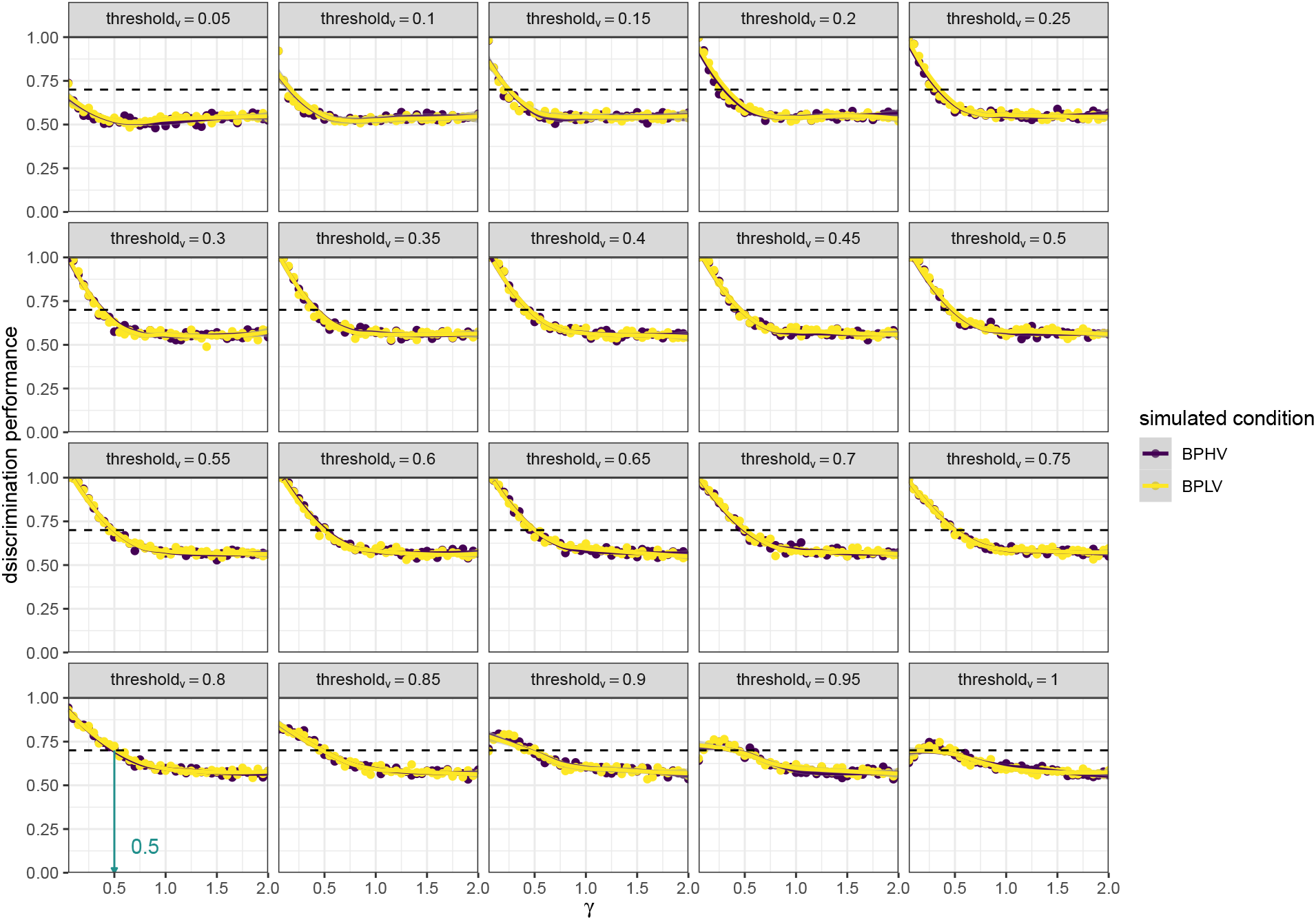
Sensitivity tests for the volume first model. Same notation as in Fig. S6. The different panels give the different values of the salience threshold that needed to be reached for one option to be preferred over the other. We set the value of the threshold for both the volume and probability dimensions to 0.8, based on the psychometric function threshold for probability (Rivalan, Winter, and Nachev 2017).

**Figure S10:**
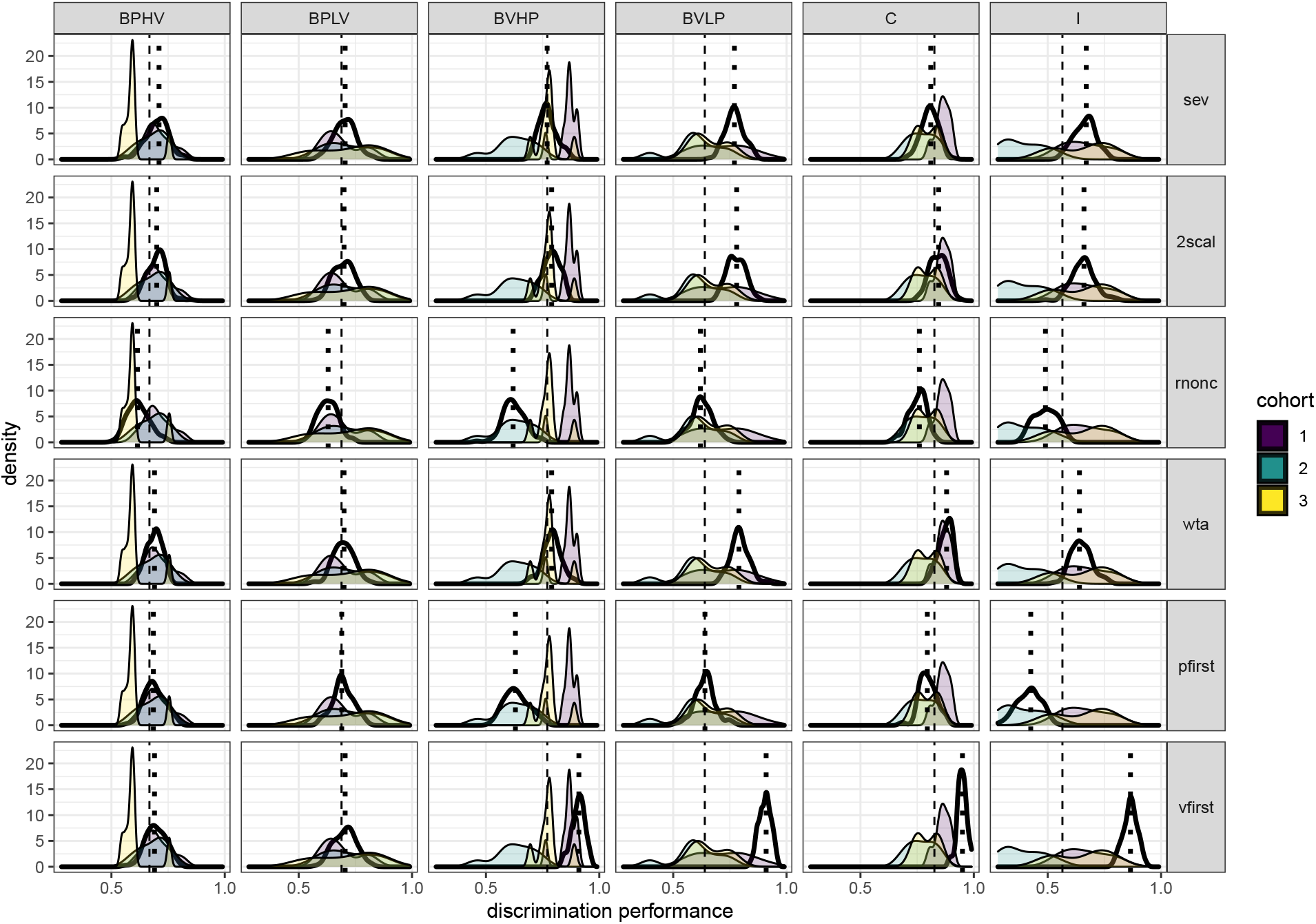
Comparison of discrimination performance in all six simulation models and in the three mouse cohorts in Experiment 1. Columns give the condition names (Fig. 2) and rows, the model number (Table 1). Empirical data from the three cohorts are represented by differently color-filled density curves from the observed discrimination performances. Simulation data are represented by an empty thick-lined density curve. The dashed line gives the median of the empirical data and the dotted line - the median of the simulated data. The discrimination performance gives the relative visitation rate of the more profitable option, or, in the incongruent condition, the option with the higher volume.

**Figure S11:**
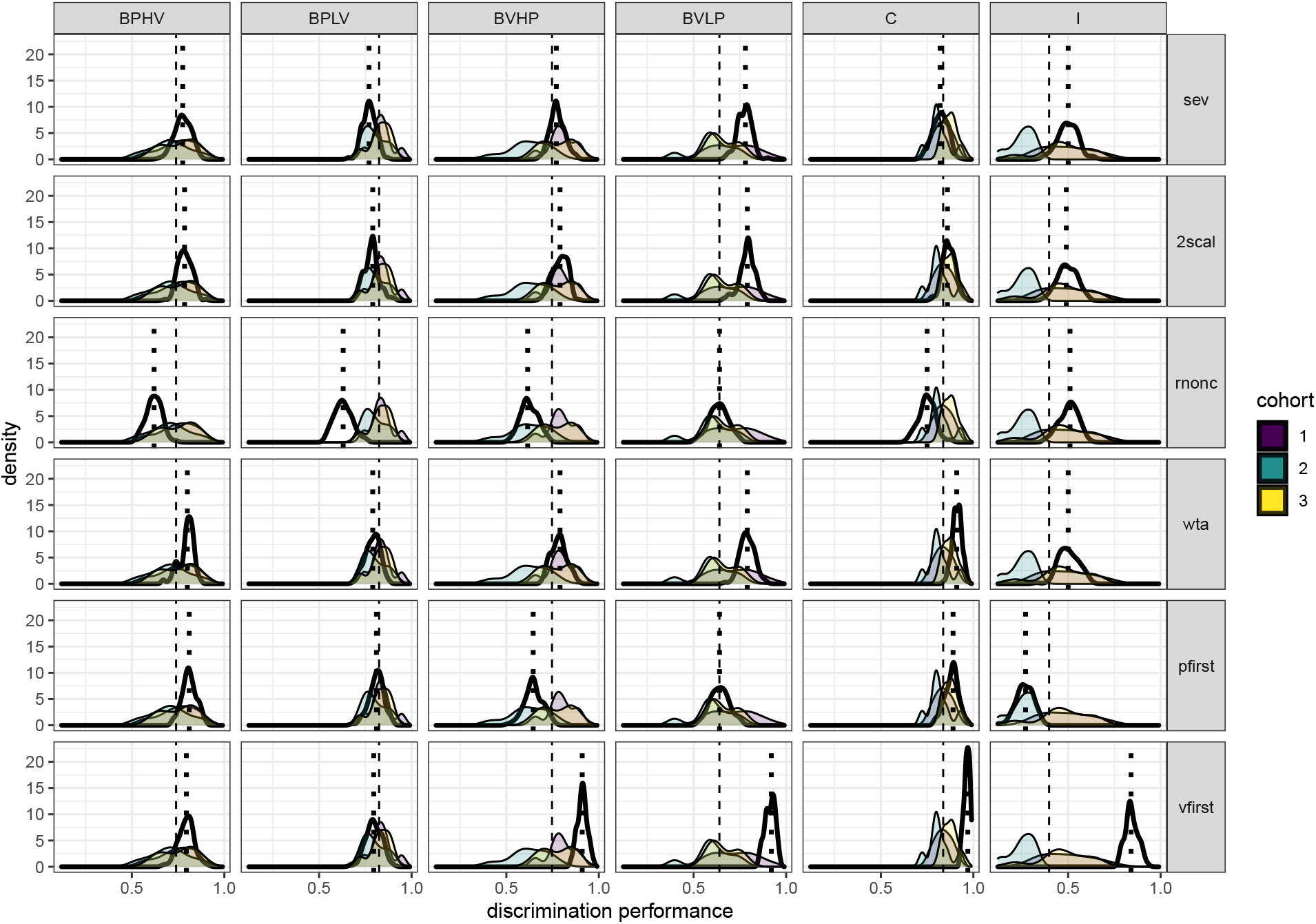
Comparison of discrimination performance in all six simulation models and in the three mouse cohorts in Experiment 2. Same notation as in Fig. S10.

**Figure S12:**
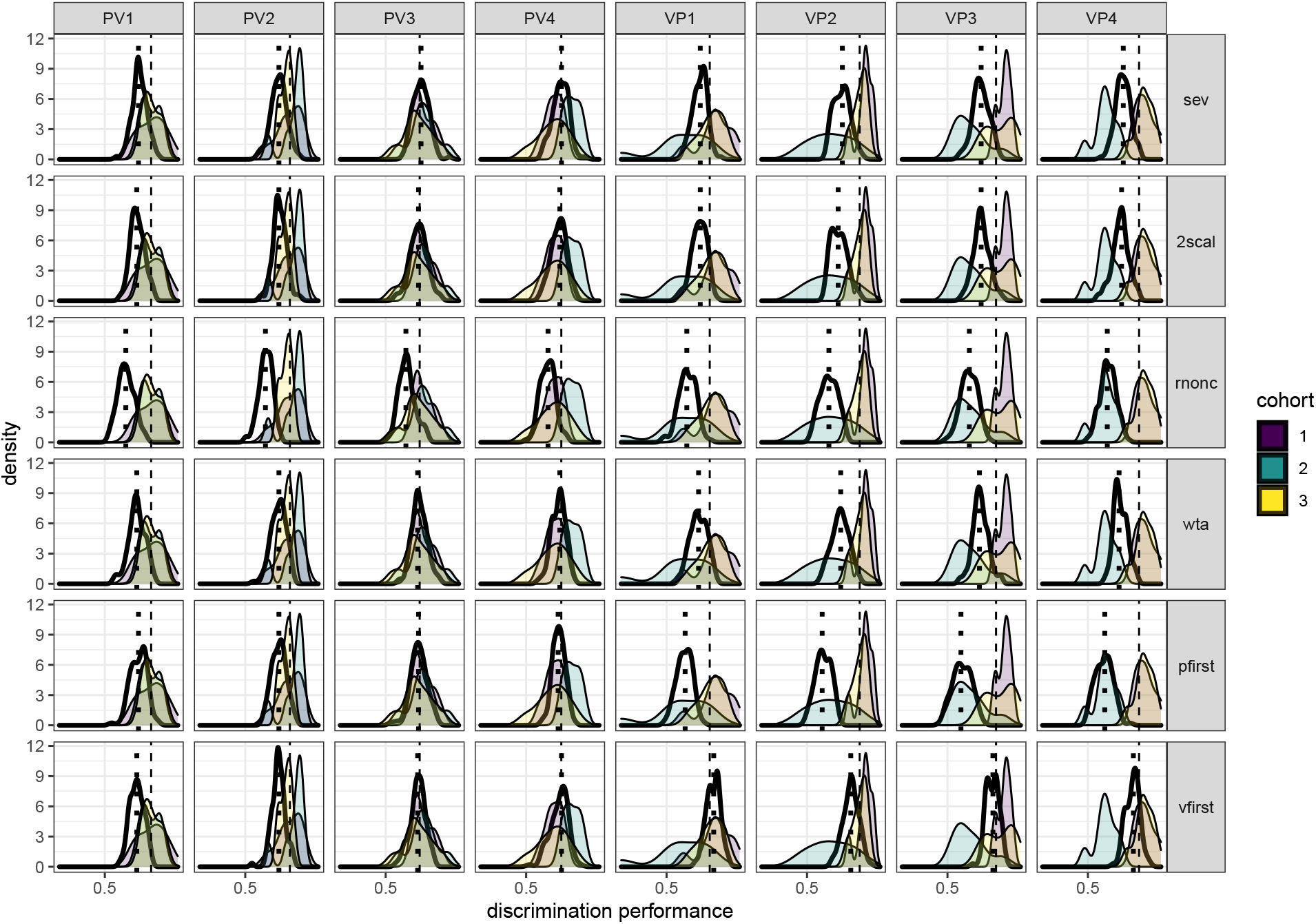
Comparison of discrimination performance in all six simulation models and in the three mouse cohorts in Experiment 3. Same notation as in Fig. S10.

**Figure S13:**
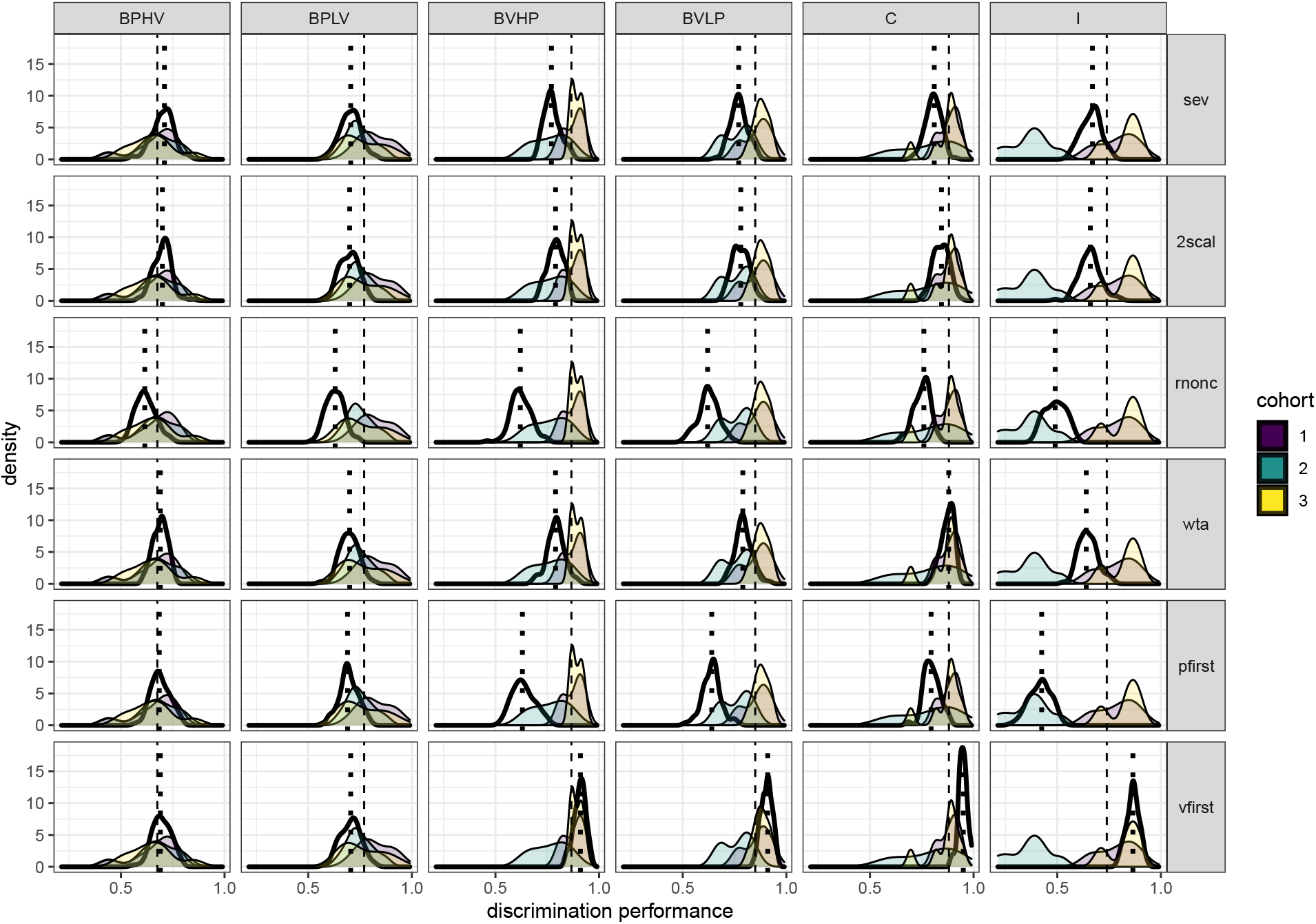
Comparison of discrimination performance in all six simulation models and in the three mouse cohorts in Experiment 4. Same notation as in Fig. S10.

**Figure S14:**
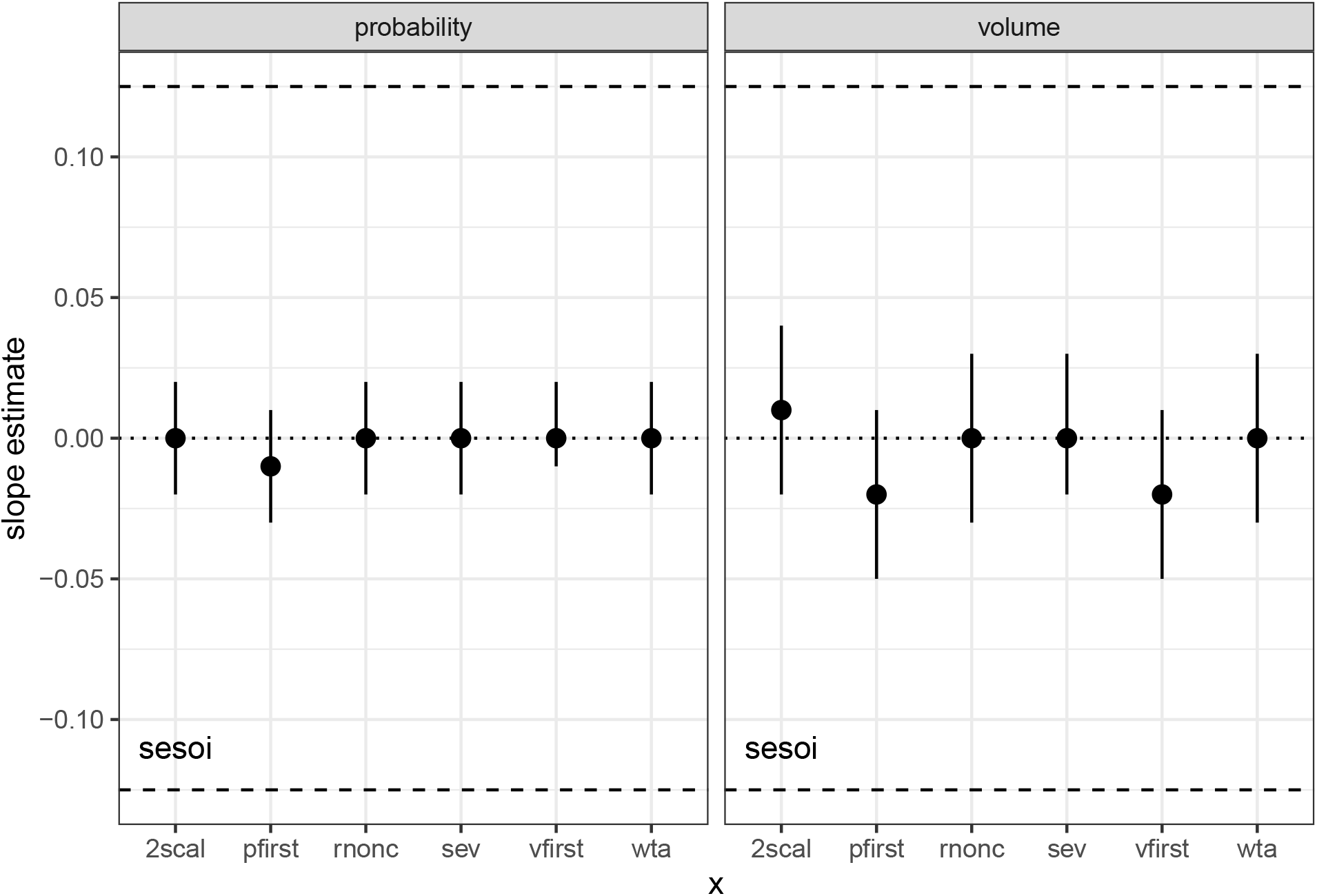
Slope estimates for the effect of the background dimension on the discrimination performance in the relevant dimension for different decision models. The two choice options always differed along the relevant dimension (either probability or volume) at a fixed relative intensity. The discrimination performance for 100 virtual mice making 100 decisions each was measured at four different levels of the background dimension. Symbols and whiskers give means and 98% confidence intervals estimated from bootstraps. The smallest effect size of interest (dashed lines) was determined to be the slope that would have resulted in a difference in discrimination performance of 0.1, from the lowest to the highest level of the background dimension. Compare to Fig. 5.

**Figure S15:**
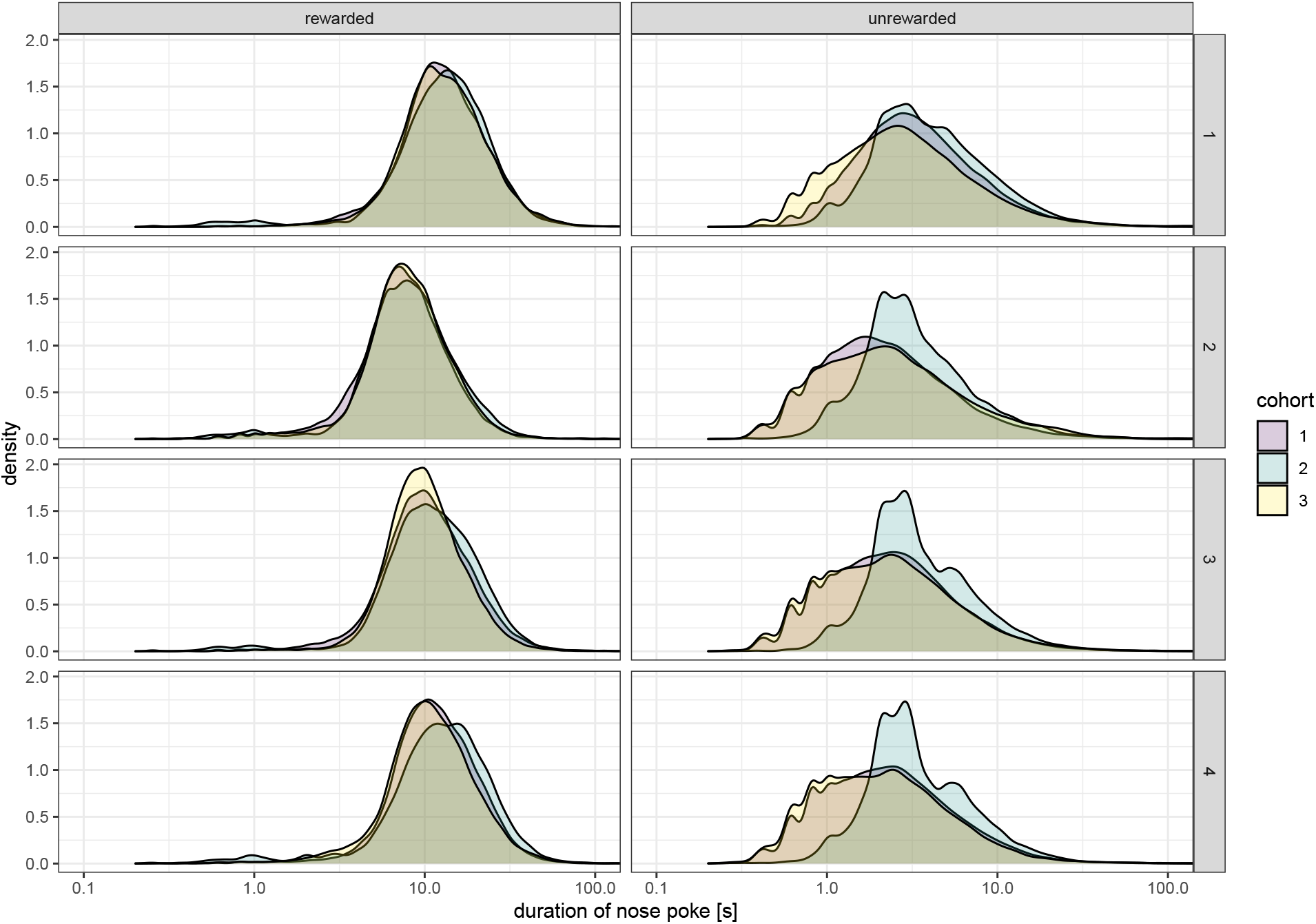
Visit durations during rewarded and unrewarded nose pokes for the three cohorts in all experiments. Columns give the status of the nose poke (rewarded or unrewarded) and rows, the experiment number (1-4). Data from the three cohorts are represented by differently color-filled density curves from the observed individual nose poke durations. Note the logarithmic scale on the abscissa.

## Declarations

### Funding

No specific funding.

### Conflicts of interest/Competing interests

YW owns PhenoSys equity.

### Ethics approval

The experiments were conducted under the supervision and with the approval of the animal welfare officer heading the animal welfare committee at Humboldt University. Experiments followed national regulations in accordance with the European Communities Council Directive 10/63/EU.

### Consent to participate

Not applicable.

### Consent for publication

Not applicable.

### Availability of data and material

All data and code are available in the Zenodo repository: https://doi.org/10.5281/zenodo.4223729.

### Code availability

All data and code are available in the Zenodo repository: https://doi.org/10.5281/zenodo.4223729.

### Authors’ contributions

V.N. Conceptualization, Methodology, Software, Formal Analysis, Data curation, Writing—original draft, Writing—review and editing, Visualization, Supervision, Project Administration.

M.R. Methodology, Writing—review and editing, Supervision.

Y.W. Resources, Methodology, Writing—review and editing, Supervision.

## Acknowledgments

We thank Miléna Brunet, Alexia Hyde, and Sabine Wintergerst for data acquisition, Katja Frei for assistance with the mice, Alexej Schatz for programming of the control software, and our colleagues of the Winter lab for a fruitful discussion. We also thank Noam Miller, Daniël Lakens, and two anonymous reviewers for their helpful comments and suggestions for improving the manuscript.

## Notes

https://doi.org/10.5281/zenodo.4590171

### Summary of Updates

General edits and added figures for clarity.

